# Fragmentation Patterns of Human Telomeric Chromatin in Plasma cfDNA

**DOI:** 10.64898/2026.05.28.728442

**Authors:** Brandon Buck, Abigail Weirich, Stephanie Eramo, Rachel Lopez, Laura K. White, Peter Kabos, Craig Forester, Srinivas Ramachandran

**Affiliations:** Department of Biochemistry and Molecular Genetics, Children’s Hospital Colorado, University of Colorado Anschutz Medical Campus, Aurora, Colorado, 80045 U.S.A; RNA Bioscience Initiative, Children’s Hospital Colorado, University of Colorado Anschutz Medical Campus, Aurora, Colorado, 80045 U.S.A; Department of Medicine, Division of Medical Oncology, Children’s Hospital Colorado, University of Colorado Anschutz Medical Campus, Aurora, Colorado, 80045 U.S.A; Department of Pediatrics, Children’s Hospital Colorado, University of Colorado Anschutz Medical Campus, Aurora, Colorado, 80045 U.S.A; Division of Pediatric Hematology/Oncology/Bone Marrow Transplant, Children’s Hospital Colorado, University of Colorado Anschutz Medical Campus, Aurora, Colorado, 80045 U.S.A

**Author notes:** These authors jointly supervised this work: Craig Forester, Srinivas Ramachandran.

## Abstract

The unique chromatin structure at telomeres protects the linear ends of chromosomes from DNA surveillance machineries. Here, we demonstrate that circulating cell-free DNA (cfDNA) from plasma can map chromatin structure at telomeres. We find that the telomeric 6-mer repeats (TTAGGG/CCCTAA) are the most abundant circulating 6-mers in cfDNA. Telomeric sequences in cfDNA contain subnucleosomal footprints distinct from the rest of the genome, arising from specific cleavages in the C-rich strand, and a nucleosome repeat length of 145 bp, markedly shorter than the ∼170 bp observed genome-wide. The abundance of cfDNA telomeric footprints decreases with age, and this decline is exacerbated by Dyskeratosis Congenita (DC), a telomere biology disorder. Promoter subnucleosome enrichment from cfDNA identifies DC-specific gene signatures that reflect disease states and show enrichment towards chromosome ends. In this work, we demonstrate that cfDNA captures telomere chromatin structure and its genome-wide impact non-invasively, including disease-specific signatures in DC.

## INTRODUCTION

Telomeres shield chromosome ends from recognition and processing by DNA repair machinery. Telomeres contain a 6-mer repetitive DNA sequence^1^ (TTAGGG, “G-rich” repeats and complementary CCCTAA, “C-rich” repeats) and form a unique chromatin structure mediated by both nucleosomes and components of the shelterin complex: TRF1^2, 3^, TRF2^2, 4, 5, 6^, and Pot1^7, 8^. This specialized chromatin architecture is essential for chromosome end protection, inhibition of the DNA damage response^9^, and regulation of telomerase activity^3^.

Circulating cell-free DNA (cfDNA) in plasma is generated by the action of endogenous nucleases on chromatin^10, 11^, and the resulting fragment length distribution directly encodes the positions and sizes of protein-DNA complexes (mainly nucleosomes) across the genome^12^. Under this framework, cfDNA is not simply a cancer biomarker but a plasma-accessible chromatin footprinting readout: any genomic region with a stable, distinct protein-DNA architecture will leave a corresponding footprint signature in cfDNA that can be recovered non-invasively from a blood draw. Nucleosomes, transcription factors, and subnucleosomal intermediates generated during transcription have all been shown to generate characteristic cfDNA footprints, and the enrichment of subnucleosomal footprints at gene promoters has previously enabled us to recover gene expression profiles from plasma^10^. Telomeres, with their unique combination of tightly phased nucleosomes and sequence-specific shelterin complexes, are precisely the kind of genomic region expected to produce a distinct and interpretable cfDNA footprint. Yet, telomeric sequences have been almost entirely overlooked in cfDNA studies, primarily due to technical challenges in aligning short sequencing reads to repetitive DNA. We reasoned that because typical sequencing reads (∼150 bp) are comparable in length to nucleosomal footprints (∼130–170 bp), an alignment-free approach that classifies reads entirely by their sequence composition could bypass this mapping problem and reveal telomeric chromatin structure directly.

Telomeres progressively shorten with each division by ∼50–150 base pairs due to incomplete end replication and telomere-specific processing^13, 14, 15, 16, 17^. With continued divisions, telomeres shorten beyond the threshold needed for loop formation, resulting in linearized ends that activate DNA damage responses^18, 19, 20^. At the cellular level, this telomere erosion enforces two proliferative barriers: senescence (mortality stage 1), a stable cell cycle arrest, and crisis (mortality stage 2), a cell death response mediated by autophagy and innate immune activation^21, 22^. Shortening of telomeres is correlated with age and occurs in a predictable manner of approximately 25 bp/year^23^. However, pathological telomere shortening leads to telomere biology disorders, such as dyskeratosis congenita (DC)^24^, caused by mutations in telomere maintenance genes^25^. DC patients have telomere lengths in the <1 to <10 percentile for their age cohort^25^ and present with altered dermatological, pulmonary, hematopoietic, and hepatic physiology^26, 27, 28^. DC patients have a >90% chance of developing bone marrow failure, have an increased risk for cancer, and there is currently no cure for this disorder^25, 29^.

Here, we show that telomeric chromatin architecture can be resolved *in vivo* through analysis of cfDNA. Telomeric cfDNA exhibited distinct footprints compared to the rest of the genome, offering insight into telomeric chromatin organization *in vivo*. We identified three footprint classes, two of which are consistent with TRF1 binding at DNA entry and exit sites and the third with phased nucleosome arrays, in agreement with prior *in vitro* biochemical reconstitutions. Notably, the abundance of telomeric footprints decreases with age and in patients with DC. We can correlate changes in promoter subnucleosome enrichment with telomere shortening in DC, suggesting possible genome-wide changes in gene expression that accompany telomere shortening in DC. In summary, we demonstrate that cfDNA provides an unexpected means to study telomere chromatin structure and connect changes in telomere state to gene expression changes that may underlie the pathological effects of telomere biology disorders.

## RESULTS

### Telomeric repeats are the most abundant circulating 6-mers

To determine the chromatin structure of a repetitive genomic region from cfDNA, we employed an alignment-free analysis framework. Our library approach separates cfDNA into individual strands before ligation^30^, ensuring that we capture nicks and the strand information of cfDNA. We restricted our analysis to the R1 read of 150 bp paired-end sequencing datasets we generated (n=16, 8 males, 8 females, age=19-75 yo, ∼2.4 billion fragments analyzed). After removing adapters, we trimmed R1 reads to 140 bp to further remove bases contributed by the adapter that might be too short to be recognized by adapter trimming software. We then extracted fragments that only contained tandem repeats of each possible 6-mer for the entire length of the R1 read. We found the telomeric repeat sequence (CCCTAA and TTAGGG) to be the most abundant 6-mers in cfDNA, 54- and 18-fold more enriched respectively than the next most abundant 6-mer repeat (**Figure 1A**). Three other repeats in the top ten most abundant 6-mer repeat protections in cfDNA correspond to telomeric variants, with only one bp edit distance from the telomeric repeat sequence (**Figure 1B**). These variants are present in the genomic reference sequence (Telomere-to-Telomere^31^ (T2T) CHM13v2.0/hs1) consistent with their occurrence in cfDNA. The difference in enrichment in telomeric repeats compared to other 6-mers is higher in cfDNA than in the genomic reference sequence (**Figure 1C**), probably because non-telomeric 6-mer repeats do not occur in stretches >140 bp for the most part (**Figure 1E**). We then analyzed the length distribution of the sequenced, and therefore protein-protected cfDNA footprints for the most abundant 6-mer repeats. Here we observed that telomeric repeats have a unique distribution of footprint lengths, with protections ranging from 60 to 140 bp (**Figure 1D**). The repeat with an edit distance of 1 from the telomeric sequence (CCCCAA) has footprints in the range of 40-60 bp, whereas the only other repeats with an abundance of at least 1 read per million have much shorter protections (**Figure 1D**). In summary, telomeric repeats are the most abundant 6-mer repeat protections in cfDNA and present a unique structural signature compared to other 6-mer repeats.

**Figure 1.**
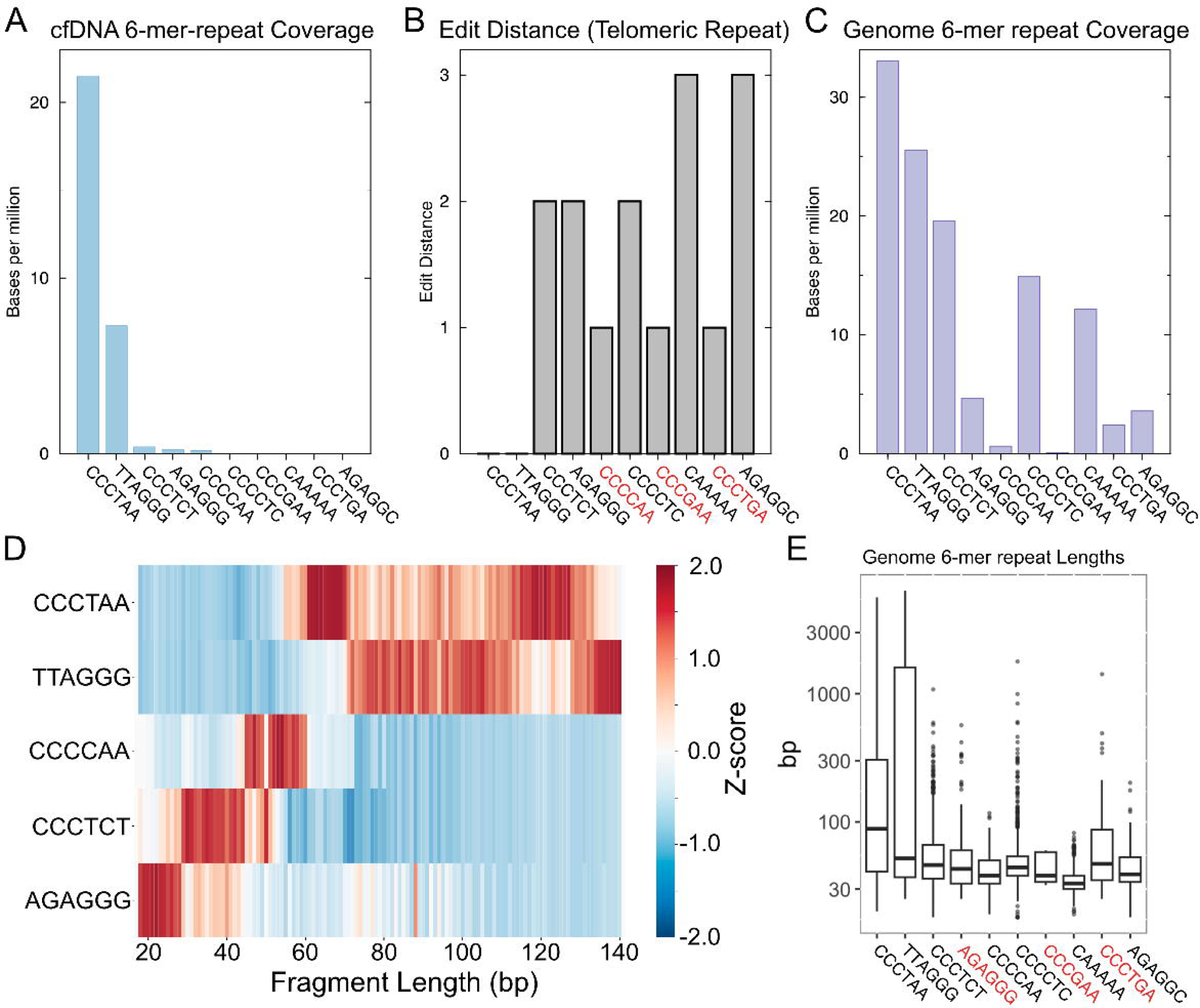
Telomeric 6-mer sequences are highly enriched and display footprints different from other 6-mer sequences in human blood plasma. **A)** Top ten 6mer sequences present in human cfDNA quantified by number of bases per million (sequences obtained from n=16 combined human cfDNA datasets). The top two 6mer sequences in the human cfDNA samples are the C-rich and G-rich telomeric repeats. **B)** Edit distance of each 6-mer shown in **(A)** to the telomeric 6-mer repeats. **C)** Genomic coverage of the 6-mer shown in **(A)**. C-rich and G-rich telomeric 6mers had the longest coverage throughout the human genome. **D)** Heatmap of the fragment length distribution of the top five 6mer sequences occurring in the human cfDNA samples. The heatmap was generated by the column-wise Z-normalization of the matrix of fragment length distributions across the top 5 6-mers. **E)** Boxplot of lengths of segments containing each of the 6-mers in the human genome. Telomere repeats had the widest range of repeat lengths, while the other 6-mer repeats ranged between 30 and 100 bp.

### Telomeric sequences contain distinct subnucleosomal footprints arising from specific cleavages in the C-rich strand

Building on the distinct subnucleosomal footprints revealed by 6-mer analysis, we next examined their genome-wide prevalence. We aligned the trimmed R1 reads to the T2T reference genome and determined the footprint size distribution at different genomic regions (**Figure 2A**). Genome-wide, we do not see an enrichment of short footprints, but rather a peak at 140 bp (see arrowhead in **Figure 2A**). A 140 bp length would represent footprints of at least 140 bp, suggesting, as expected, that most of the genome is enriched for nucleosomal footprints. Next, we analyzed the fragment length distribution of α-satellite repeat sequences found in the centromeric and pericentromeric regions, which contain constitutive heterochromatin. α-satellite repeat sequences featured fragment length distributions similar to the genome-wide distribution, with the main enrichment observed at 140 bp. Finally, we analyzed the fragment length distribution at CTCF binding sites, as defined by ChIP-seq^32^, in GM12878, a lymphoblastoid cell line. CTCF is a constitutively expressed transcription factor with a long residence time on DNA. We have previously demonstrated that CTCF can generate short footprints in cfDNA that correlate with binding strength in the tissue of origin of the cfDNA^32^. Even CTCF binding sites, which have known enrichment of footprints between 70 and 100 bp, are not as highly enriched for short footprints as G-rich and C-rich telomeric repeat sequences (**Figure 2A**). This finding leads to two possible hypotheses: either sequence-dependent cleavages lead to the observed footprints, or specific chromatin structures formed on telomeric sequences might lead to the observed footprint populations. These short footprints are not indicative of sequence-driven cleavages due to nuclease bias: with 6*-*mer repeats, we would expect sequence-driven cleavages every six base pairs, not distinct footprint populations. Instead, binding of specific protein structures is more consistent with these footprint populations. This analysis suggests that telomeres have a chromatin structure distinct from the rest of the genome and are characterized by short footprints that remain preserved in plasma cfDNA.

**Figure 2.**
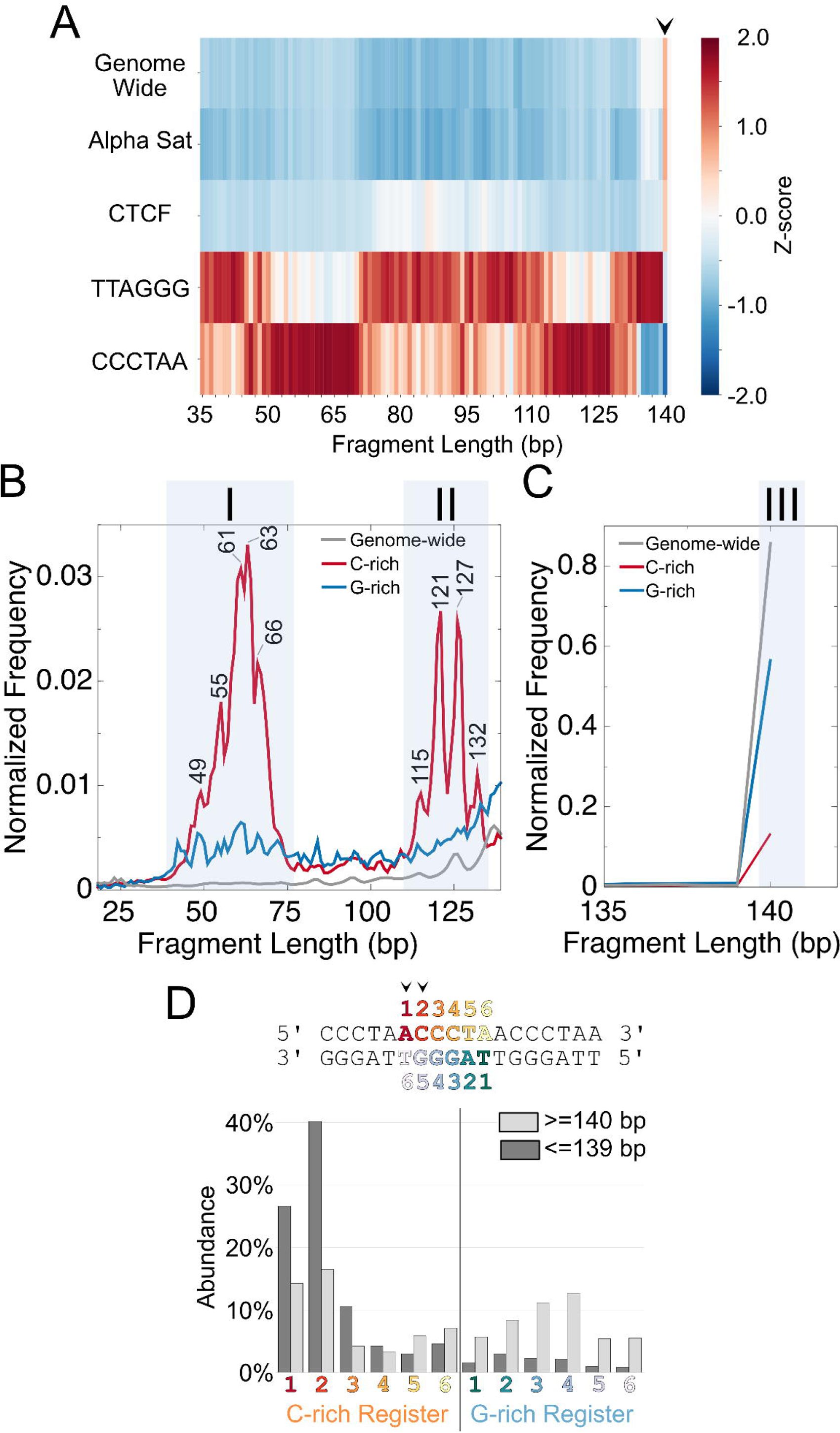
Telomeric sequences contain distinct subnucleosomal footprints arising from specific cleavages in the C-rich strand. **A**) Heatmap of the fragment length distribution genome-wide, at α-satellite sequences, CTCF binding sites, and G- and C-rich telomeric sequences. The heatmap was generated by the column-wise Z-normalization of the matrix of fragment length distributions across the 5 sequence classes analyzed. Arrowhead points to >=140 bp footprints. **B**) Fragment length distribution of the telomeric cfDNA in the range 18-135 bp. Two distinct classes of footprints are observed for C-rich sequences, which are consistent with subnucleosomal (class I) and short nucleosomal (class II) complexes. **C**) Same as (**B**) for fragments in the range 135-140 bp, showing apparent nucleosomal footprints for C-rich and G-rich telomeric sequences and genome-wide cfDNA (class III). **D**) Abundance of the register of the telomeric repeat (top), where 5’ end cleavages occur on cfDNA, shown as a bar plot (bottom). We observe a clear enrichment for positions 1 and 2 in the C-rich strand for fragments <140 bp (pointed by arrowheads on the sequence). For all analyses in this figure, sequences obtained from n=16 combined human cfDNA datasets were used.

In our approach, we are able to observe differences in footprint sizes between C-rich and G-rich telomeric sequences because our libraries are stranded and capture nicked DNA as well. In the G-rich telomeric sequences, we observed an increased abundance of shorter footprints, ranging from 40 to 140 base pairs, compared to the genome-wide footprint size distribution (**Figure 2B**). In contrast, in C-rich telomeric sequences, we observe three distinct populations of footprints. The first population consists of six peaks in the 45-75 bp range, which we interpret as subnucleosomal footprints. The second population of footprints consists of four peaks in the 115-135 bp range, which we interpret as short nucleosomal footprints due to nucleosome unwrapping^10^. For both populations of footprints, a subset of peaks is separated by six bp (**Figure 2B**). To ask if we are able to observe the shorter footprints because our libraries are stranded and capture nicked DNA, we analyzed published cfDNA datasets generated by standard double-stranded libraries^33^. We observe no enrichment for footprints <139 bp, confirming that these footprints are lost during with double-stranded library protocols (**Supplementary Figure S1A**). Finally, the third population is characterized by footprints of at least 140 bp, which we interpret as nucleosomal footprints. These are observed with standard double-stranded libraries as well (**Supplementary Figure S1B**). The nucleosomal footprints have higher enrichment genome-wide compared to G-rich and C-rich telomeric sequences, and G-rich telomeric sequences have higher enrichment than C-rich telomeric sequences (**Figure 2C**). The two specific populations of footprints observed only in the C-rich strand suggest that protein binding creates specific accessibility uniquely in the C-rich strand.

TRF1 and TRF2 bind telomeric repeats in a sequence-specific manner. If C-rich footprints arise due to the binding of these proteins, the footprints are expected to have a specific register consistent with the mode of binding of TRF1 and TRF2 to the telomeric repeats via their Myb domains (TRF1 and TRF2 identically bind telomeric repeats)^34^. To ask if specific 5’ registers are enriched in the footprints, we calculated the 5’ register for G- and C-rich subnucleosomal and short nucleosome footprints (<=139 bp) and nucleosomal footprints (>=140 bp). The footprints with sizes >=140 bp do not have a preferred register (**Figure 2D**). The lack of a preferred register for footprints with sizes >=140 bp suggests that: i) nucleosomes may not have a preferred position on the telomeric repeats, and ii) the footprints are not simply arising due to nuclease cleavage preference^35^ of a specific register of the telomeric repeat. In contrast, the footprints with sizes <=139 bp display a striking enrichment of 5’ACCCTA and 5’CCCTAA registers (**Figure 2D**). In the crystal structures of TRF1 and TRF2 Myb domains, these positions on the telomeric repeat sequence are at the edge of the protein-DNA interface, suggesting that these may be the positions accessible to the nuclease right outside the protein binding site^34^. To further rule out nuclease bias driving the preferred registers for telomeric fragments <139 bp, we analyzed single nucleotide and dinucleotide frequency at 5’ and 3’ ends of both telomeric repeat-containing sequences and all sequenced fragments. We observe a preference for C on the 5’ end (consistent with “CCCTAA” register) only in <=139 bp fragments for the telomeric sequences. Genome-wide, there is no difference in sequence preference between <=139 bp fragments and >=140 bp fragments (**Supplementary Figure S1C**). Similarly, the 3’ end single nucleotide, and both 5’ and 3’ end dinucleotides show that the preferred registers for telomeric sequences do not arise from nuclease bias, and specific protein protections are a more parsimonious explanation (**Supplementary Figure S1D-F**).

A recent cryo-EM structure of shelterin subcomplex reconstituted with the nucleosome suggested partial unwrapping of the nucleosome to accommodate TRF1 binding at the nucleosome entry/exit sites^36^. The extent of unwrapping was determined to be 1.5 turns on either side of the nucleosome, corresponding to footprints of 117 and 132 bp. These sizes match well with our short nucleosomal footprints (population II). Furthermore, if nucleosome wrapping is constrained based on TRF1 binding at the edge of the nucleosome, these unwrapped nucleosomes are expected to have a specific positioning on the telomeric sequence due to TRF1’s binding register. As short nucleosomal footprints are included in the footprints of size <139 bp, they are enriched for the 5’ACCCTA and 5’CCCTAA registers as well, consistent with these footprints arising from unwrapped nucleosomes adjacent to a bound TRF1. In summary, telomeric repeats have two short footprint classes that are not observed in other parts of the genome, and these short footprints are consistent with the unique protein and nucleosome complexes expected to be formed at telomeres. These results suggest that cfDNA provides a novel approach to understanding telomeric chromatin structure.

### Long-read sequencing identifies nucleosomal repeat length of telomeric cfDNA

Because 150 bp paired-end sequencing (PE150) restricts the maximum telomeric footprint sizes that we can accurately measure to 139 base pairs, we used long-read sequencing on the Oxford Nanopore Technology (ONT) platform to identify footprint lengths beyond 139 base pairs. To enrich for telomeric sequences and accurately measure footprint lengths, we designed a library preparation approach that utilized single-strand adapter ligation before proceeding to nanopore library preparation (**Figure 3A**). First, we could determine the originating strand because ligation was performed on single strands. The Illumina P5 adapter was ligated to the 5’ end of single-stranded DNA and the Illumina P7 adapter was ligated to the 3’ end enabling strand identification post-sequencing. Second, by ligating Illumina P5 and P7 adapters to cfDNA fragments after separating them into single strands, we create unique junctions between the repetitive DNA ends and the adapters. This allows us to amplify repeat sequences while preserving their ends. We enriched for telomeric cfDNA footprints using primers that selectively amplified either the C-rich strand or the G-rich strand. As an alternative, we also used telomeric 6-mers as splint sequences for the splinted ligation step of single-strand ligation, which theoretically enhances the ligation of adapters to telomeric sequences (**Supplementary Figure S2A**).

**Figure 3.**
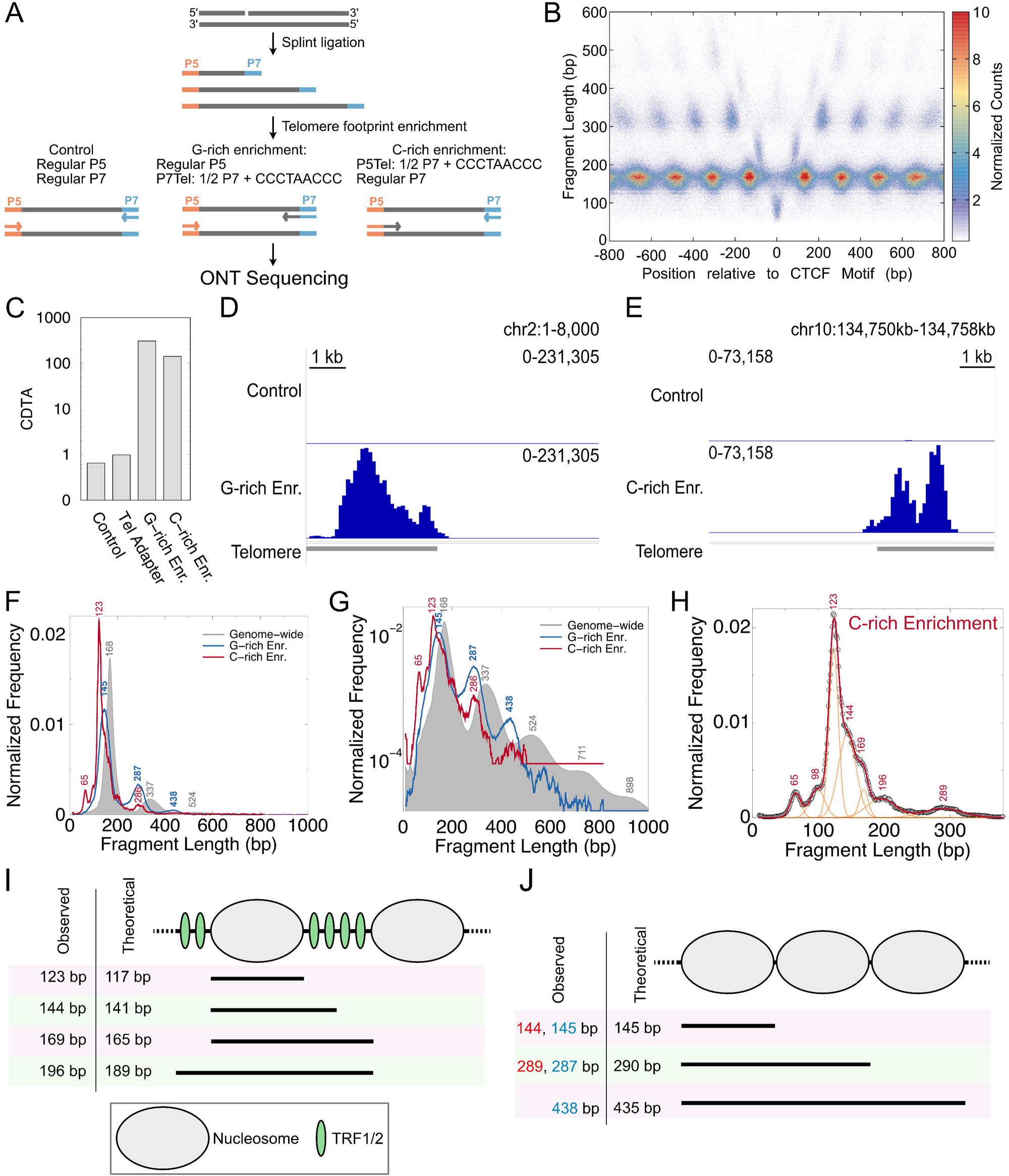
Long-read sequencing identifies nucleosomal repeat length of telomeric cfDNA. **A**) Schematic of library strategy used to enrich for telomeric sequences from cfDNA while preserving fragment ends. **B**) Fragment midpoint vs. length plot (“V plot”) at CTCF motifs that are enriched for short footprints in cfDNA. **C**) cfDNA telomeric abundance (CDTA) plotted for different library enrichment strategies. **D**) Genome browser snapshot showing a representative telomere for non-enriched and G-rich strand-enriched libraries. **E**) Same as (**D**) for C-rich strand enriched libraries. **F**) Fragment length distribution for all fragments (“Genome-wide”), fragments with only G-rich telomeric repeat sequences, and fragments with only C-rich telomeric repeat sequences. **G**) Same as (**F**), but the normalized frequency is plotted on a log10 scale to highlight lower abundance peaks from the fragment length distribution. **H**) The fragment length distribution of C-rich telomeric repeats fitted with 7 Gaussian distributions is shown. The original fragment length distribution is shown with circles, the sum of the 7 Gaussian distributions is shown with a red line, and the individual distributions are shown with orange dashed lines. The mean value for each Gaussian distribution is indicated on the plot. **I**) Model for footprints generated with TRF1/2 binding adjacent to nucleosomes. **J**) Model for footprints generated by nucleosome arrays formed on telomeric repeats.

We performed five total ONT sequencing runs: two runs of control sequencing without any enrichment, one run of G-rich PCR enrichment, one run of C-rich PCR enrichment, and one run where libraries were generated using telomeric splint during ligation. We used a custom adapter trimming code with high stringency to identify sequencing reads where both P5 and P7 adapters could be trimmed. This allowed us to obtain footprint sizes from ∼50 bp to ∼1 kb from cfDNA. To ensure we were faithfully capturing cfDNA footprint sizes, we generated a fragment length vs. midpoint plot (“V-plot”^37^) at CTCF binding sites that are enriched for short cfDNA footprints based on the Illumina sequencing datasets. To generate the V-plot, we combined aligned reads from all five sequencing runs and selected fragments whose centers aligned within 800 bp of CTCF binding sites. For a protein-protected region in the genome, the V-plot would show a “V” with the vertex at 0 on the x-axis and at the length of the minimal footprint on the y-axis of the V-plot. For CTCF, we observe a clear central “V” from the long-read sequencing data at sites expected to have a strong footprint based on short-read sequencing (**Figure 3B**, annotated V-plot in **Supplementary Figure S2C**). At CTCF binding sites where footprints are likely not expected based on short-read sequencing, we do not observe a “V” or any distinct patterns in the V-plot (**Supplementary Figure S2B**). Thus, the V-plot from nanopore sequenced cfDNA fragments revealed that our library strategy faithfully preserved cfDNA footprint lengths and positions, allowing us to obtain robust structural signatures. To confirm that our nanopore pipeline recapitulates established cfDNA footprint signatures, we also generated V-plots at CTCF binding sites from our short-read Illumina cfDNA datasets, where CTCF footprints are well-characterized from our prior work (**Supplementary Figures S2D, E**). The concordance between the Illumina and nanopore V-plots at the same CTCF sites confirms that the nanopore adapter-trimming pipeline faithfully preserves fragment length information, validating its use for measuring the novel footprint sizes observed at telomeric sequences.

We also observe the phased positioning of nucleosomes upstream and downstream of the CTCF binding site (increased density at fragment lengths of about 150 bp). Additionally, the longer reads allow us to observe positioned dinucleosomes (fragment lengths slightly above 300 bp) and trinucleosomes (fragment lengths around 500 bp). Finally, we also observe specific signals for CTCF co-bound with either an upstream nucleosome, a downstream nucleosome, or both (**Figure 3B**, annotated V-plot in **Supplementary Figure S2C**). Thus, we obtain a complete picture of all possible footprints, revealing co-occupancies of nucleosomes and CTCF. In summary, the V-plot demonstrates that our approach, which combines adapter ligation and nanopore sequencing, yields high-resolution, long footprints from cfDNA.

We next identified reads that are entirely made of telomeric repeats (which we will denote as “telomeric reads”), similar to our earlier analysis (**Figure 2**). We quantified the enrichment of telomeric reads as cfDNA Telomeric Abundance (CDTA, see **Methods**). The approach of using telomeric sequences in the splint part of the splinted adapter (“Tel Adapter”) resulted in only a modest increase in CDTA (**Figure 3C**). However, using primers that were hybrids of P5 or P7 adapter sequence and telomeric sequence for PCR enrichment of libraries resulted in a greater than 100-fold increase in CDTA for the enrichment of both the G-rich strand and the C-rich strand (**Figure 3C**). We observed telomere-specific enrichment in genome browser snapshots comparing the unenriched cfDNA long-read sequencing dataset to either the G-rich-enriched (**Figure 3D**) or the C-rich-enriched datasets (**Figure 3E**).

To generate fragment length distributions, we combined reads from all five datasets and filtered separately for G-rich telomeric reads and C-rich telomeric reads. We observed peaks in the fragment length distribution that were unique to G-rich and C-rich telomeric reads compared to the peaks observed in the length distribution of all sequenced fragments (**Figure 3F**). Genome-wide, the mononucleosomal peak was observed at 168 bp, as expected for cfDNA^38^. In contrast, the G-rich telomeric reads had a shorter mononucleosomal footprint at 145 bp. The C-rich telomeric reads had an even shorter mononucleosomal footprint at 123 bp, corresponding predominantly to the “short nucleosome” expected from partial unwrapping due to TRF1 binding. To better explore footprints larger than mononucleosomes, we plotted the y-axis of the fragment length distribution on a logarithmic scale. With a logarithmic scale for the normalized frequency of fragments, the di-, tri-, tetra-, and even penta-nucleosomal footprints were observed genome-wide (**Figure 3G**). The dinucleosome footprint peak suggested a repeat length of ∼167 bp genome-wide, whereas the repeat length for tri-nucleosomes and beyond was observed to be ∼187 bp. Notably, 167 bp and 187 bp fragments are characteristic of footprints from cfDNA, corresponding to chromatosomes^38^. In the log-scale plot, the G-rich telomeric reads displayed a clear repeat length of ∼145 bp (**Figure 3G**). The di- and tri-nucleosome peaks for the C-rich reads also displayed a similar repeat length. Notably, the multi-nucleosomal footprints from telomeres were significantly shorter compared to the genome-wide distribution, consistent with a shorter repeat length observed in *in vitro*, cellular, and animal studies^39, 40, 41^. For fragments less than 300 bp, the C-rich reads featured a rich multimodal distribution (**Figure 3H**). We were able to robustly fit seven Gaussian distributions whose sum could explain the observed fragment length distribution of C-rich reads.

From the Gaussian fits, the pattern among a subset of the mean values suggested that the combination of two minimal footprints could explain the observed peaks (**Figure 3I**). Based on cryo-EM structures, TRF1 binding at the entry and exit sites of a nucleosome would unwrap 1.5 turns of DNA at the entry and exit sites, resulting in a nucleosomal footprint of 117 bp. Possible TRF1/2 footprints of 24 bp on either side of this “short nucleosome” explain four out of the seven peaks observed in the C-rich fragment length distribution (**Figure 3I**). The peaks common to both G-rich and C-rich fragment length distributions, namely the ∼145 bp and ∼290 bp peaks, suggest a nucleosome array with a repeat length of ∼145 bp, with even the tri-nucleosome footprint observed for the G-rich reads (**Figures 3G, J**). Importantly, the di- and tri-nucleosome footprints cannot be explained by the combination of TRF1/2 and nucleosomal binding, suggesting the existence of two categories of chromatin structures on telomeric sequences (**Figure 3I, J**). In summary, using long-read sequencing of telomeric cfDNA footprints, we have uncovered the telomeric nucleosome repeat length in humans, and the footprint ensemble suggests two possible models of telomeric chromatin landscapes.

### Dyskeratosis Congenita decreases telomeric cfDNA abundance and alters subtelomeric chromatin accessibility

The footprint distribution of telomeric sequences in cfDNA suggested that cfDNA might reflect the protein-bound state of telomeres in humans. Since telomere state is as important for end protection as telomere length^42^, we reasoned that cfDNA Telomeric Abundance (CDTA) might reflect the physiological state of the donor. Hence, we next determined the abundance of telomeric footprints as a function of age. We calculated CDTA for each donor, dividing donors into two groups based on age: young donors (19-31 years old, comprising four males and four females) and old donors (58-75 years old, comprising four males and four females). We also included cfDNA from 3 pediatric donors (4-11 years old comprising one male and two females). We then compared the distribution of CDTA between the three groups. CDTA decreases significantly with age in our cohort (**Figure 4A**).

**Figure 4.**
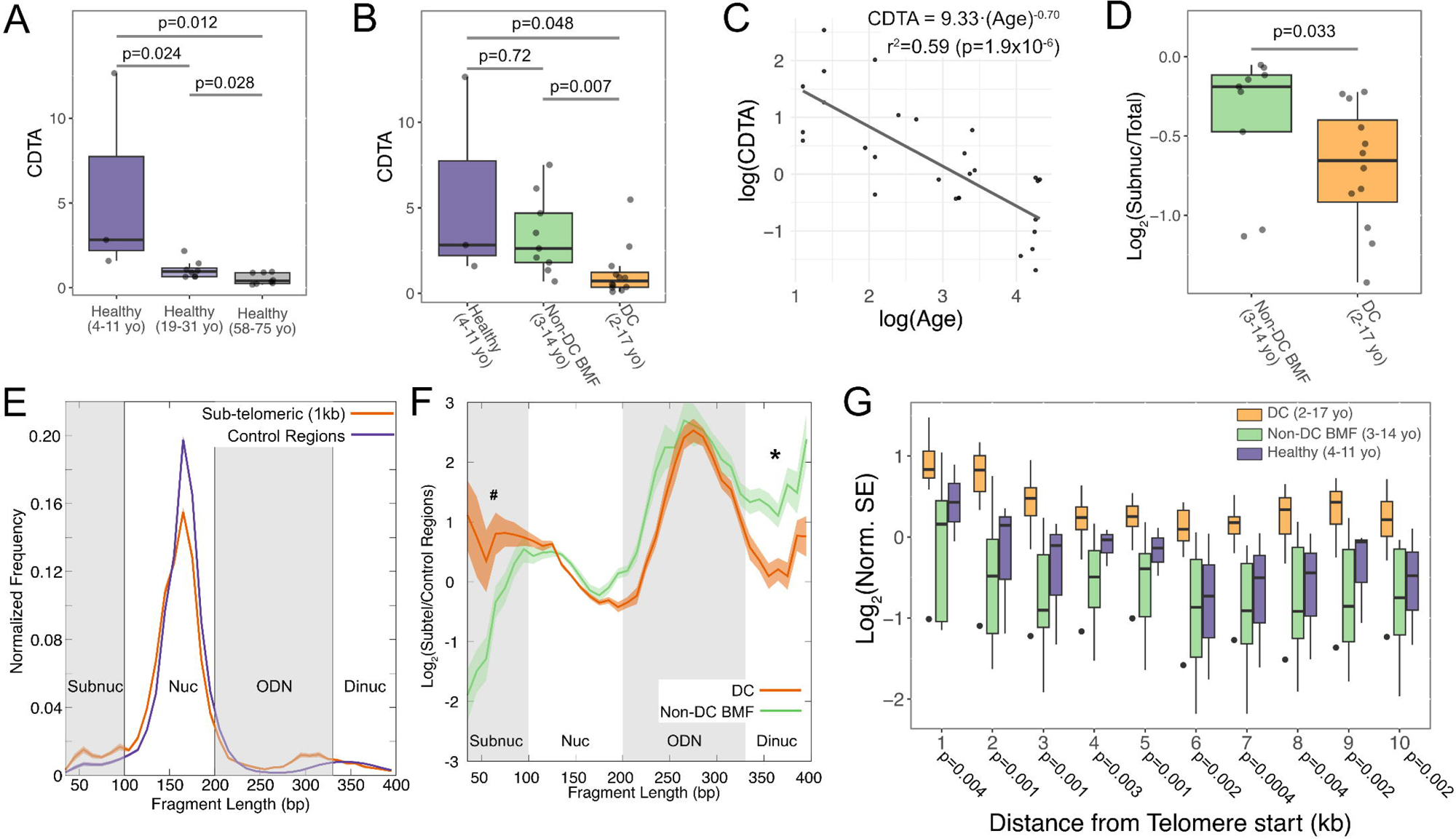
Dyskeratosis Congenita is characterized by decreased telomeric cfDNA abundance and altered subtelomeric chromatin accessibility. **A**) Cell-free DNA telomeric abundance (CDTA) was quantified for healthy pediatric (4-11 years old, n=3), healthy young (19-31 years old, n=8), and healthy old (58-75 years old, n=8) donors. **B**) Same as (**A**) for healthy pediatric donors (4-11 years old, n=3), donors with Non-DC bone marrow failure (Non-DC BMF, 3-14 years old, n=9), and Dyskeratosis Congenita (DC) donors (2-17 years old, n=12). P-values were calculated using the Wilcoxon Rank Sum test. **C**) Log-log plot of CDTA vs. Age for all donors except DC (n=28) shows a significant negative correlation. Linear model: CDTA = a.log(Age)^b^; a=9.33, 95% CI [4.5, 19.3]; b=-0.70, 95% CI [-0.93,-0.46]. P-value was calculated using F-test. F-statistic (1, 26) = 37.15 **D**) The log2 values of the fractions of telomeric cfDNA subnucleosomal and short nucleosomal footprints (45 – 75 bp) and (115 – 135 bp) were quantified for non-DC BMF and DC donors, and the distribution across the cohorts is shown as a boxplot. P-values were calculated using the Wilcoxon Rank Sum test. **E**) The length distribution of fragments mapping to the first kb upstream of the telomere (Sub-telomeric 1 kb) and fragments mapping to control regions of the human genome (corresponding to gene deserts) are plotted. The regions of the fragment length distributions corresponding to subnucleosomes (“Subnuc”), nucleosomes (“Nuc”), overlapping dinucleosomes (“ODN”), and dinucleosomes (“Dinuc”) are indicated with gray/white shaded boxes. The mean fragment length distributions across the cohort are plotted, with a shaded region indicating the standard error of the mean (SEM) around the lines. **F**) The average log2 ratio of the fragment length distributions across Non-DC BMF and DC cohorts is plotted as lines, with a shaded region around the lines indicating S.E.M. Different footprint classes are indicated with gray and white shaded boxes similar to (**C**). “**#**” marks the subnucleosome region and “**\***” marks the dinucleosome region with higher normalized enrichment of fragments in DC cohort compared to the Non-DC BMF cohort. **G**) The distribution of subtelomeric subnucleosome enrichment (SE) normalized to SE at control regions across pediatric healthy, Non-DC BMF, and DC cohorts in 1 kb windows upstream of telomeres is shown as boxplots. P-values were calculated using the Wilcoxon Rank Sum test comparing Non-DC BMF and DC cohorts.

If CDTA reflects the protein-bound state of telomeres, CDTA is expected to decrease even more dramatically in patients with telomere biology disorders. To test this hypothesis, we obtained plasma samples from patients with the telomere biology disorder, dyskeratosis congenita (DC, n=12), aged 2-17 years, comprising eight males and four females **(Table 1)**. All patients exhibited signs of cytopenias and varying degrees of bone marrow hypocellularity but had not previously undergone transplantation. DC is a rare disorder (prevalence of 1:1,000,000 Ref.^43^) characterized by pleiotropic developmental abnormalities, a heightened risk of bone marrow failure and cancer, and telomere lengths commonly <1^st^ percentile per age caused by mutations in telomeric maintenance genes. The DC patients in our cohort presented with pathogenic lesions that affected RTEL1 (n = 5), DKC1 (n = 2), and TERT (n = 5) (**Table 1**). TERT is the telomerase reverse transcriptase essential for extending telomeres in stem and progenitor cells^44^. DKC1 is necessary for maintaining human telomerase RNA levels^45^. Mutations in DKC1 result in a dramatic decrease in human telomerase RNA levels, without affecting the levels of other telomerase ribonucleoprotein components^24^. RTEL1 is a helicase essential for telomere maintenance^46^. We observed that the CDTA measurements across the DC cohort were dramatically lower compared to those in the pediatric healthy cohort (**Figure 4B**).

**Table 1.** Dyskeratosis congenita patient characteristics.

| ID | Age | Sex | Telomere Length by Q-FISH |  |  |  | Telomere age-matched %ile |  |  |  | Mutation |
| --- | --- | --- | --- | --- | --- | --- | --- | --- | --- | --- | --- |
|  |  |  | Lymph<sup>1</sup> | Gran<sup>2</sup> | B<sup>3</sup> | CD45RA+<sup>4</sup> | Lymph<sup>1</sup> | Gran<sup>2</sup> | B<sup>3</sup> | CD45RA+<sup>4</sup> |  |
| BMF16 | 4 | M | 4.8kb | 8.8kb | 7.2kb | 7.5kb | <1% | 1-10% | <1% | <1% | RTEL1 (c.2761G>C), TERT (c.2411G>T)<br>Compound Heterozygote |
| BMF24 | 17 | M | 6.0kb | 5.2kb | 6.1kb | 6.3kb | <1% | <1% | <1% | <1% | Heterozygous RTEL1 (c.2939T>G) |
| BMF28 | 2 | F | 4.4kb | 4.4kb | NSQ | 4.5kb | <1% | <1% | <1% | <1% | RTEL1_Compound_Heterozygous<br>(c.3791G>A, c.2941C>T) |
| BMF34 | 16 | M | 6.1kb | 7.2kb | 5.0kb | 6.6kb | <1% | 1-10% | 1-10% | 1-10% | 5p15.33-5p15.31 deletion (involves TERT);<br>5q33.2-5q35.3 microduplication (involves<br>NPM1 and NHP2) |
| BMF36 | 14 | M | 4.37kb | 4.22kb |  |  | <1% | <1% |  |  | DKC1_Hemizygous (c.915+10G>A) |
| BMF42 | 8 | M | 6.2kb | 5.2kb | 6.1kb | 6.5kb | <1% | <1% | <1% | <1% | RTEL1, Heterozygous, c.2139C>A<br>(p.His713Gln) |
| BMF57 | 11 | F | 6.3kb | 6.1kb | NSQ | 6.6kb | <1% | <1% | NSQ | <1% | de novo 5p15.33 deletion including full TERT<br>mutation.↗↑ Telomere <1% |
| BMF64 | 7 | M | 6.4kb | 5.0kb | 6.5kb | 6.8kb | <1% | <1% | <1% | <1% | RTEL1, c.3028C>T, p.Arg1010Ter,<br>heterozygous, pathogenic, maternally inherited |
| BMF69 | 15 | F | 3.8kb | 4.1kb | 5.2kb | 4.7kb | <1% | <1% | <1% | <1% | heterozygous for the familial pathogenic TERT<br>variant (c.1891C>T, p.Arg631Trp) |
| BMF70 | 9 | F | 5.2kb | 5.5kb | 5.9kb | 5.7kb | <1% | <1% | <1% | <1% | heterozygous for the familial pathogenic TERT<br>variant (c.1891C>T, p.Arg631Trp) |
| BMF71 | 6 | M | 5.5kb | 5.7kb | 6.1kb | 6.0kb | <1% | <1% | <1% | <1% | heterozygous for the familial pathogenic TERT<br>variant (c.1891C>T, p.Arg631Trp) |
| BMF73 | 6 | M | 5.8kb | 5.0kb | 5.8kb | 6.1kb | <1% | <1% | <1% | <1% | hemizygous sequence DKC1 variant,<br>c.473G>A p.(Arg158Gln)↗↑ |

<sup>1</sup>Lymphocytes, <sup>2</sup>Granulocytes, <sup>3</sup>B cells, <sup>4</sup>CD45RA+ T cells; NSQ = Insufficient Quantity for Reliable Analysis.

To ask if the decrease in CDTA was due to mutations in genes associated with telomere maintenance rather than a general consequence of stress or bone marrow failure, we next sequenced samples obtained from donors with other rare bone marrow failure disorders. In these disorders, the inherited mutations do not directly cause drastic telomere shortening, and patients have longer telomeres compared to patients with DC^47^. We sequenced cfDNA from donors with Fanconi anemia (n=3, two males and one female, aged 3-8 years old), Diamond-Blackfan anemia (n=2, one male and one female, aged 3 years old), and Shwachman-Diamond Syndrome (n=4, four males, aged 4-14 years old). We denote this cohort as “Non-DC BMF” with a total of 9 samples (3-14 years old, **Table 2**). CDTA for Non-DC BMF cohort is not significantly different from the pediatric healthy cohort, and greater than DC cohort with high statistical significance (**Figure 4B**). To assess technical variation in CDTA, we analyzed multiple samples collected from same donors. For one of the DC donors (BMF16), an extra tube of plasma was available, and for two of the DC donors (BMF16, and BMF36), longitudinal samples taken 13 and 25 months since the first samples were available. There was no disease progression at the later collection dates. CDTA measurements were stable across the longitudinal and repeat samples, staying well within the range of values observed for the DC cohort, confirming technical reproducibility of CDTA measurements from plasma (**Supplementary Figure S3B**).

**Table 2.** Non-Dyskeratosis congenita bone marrow failure patient characteristics.

| ID | Age | Sex | Condition | Mutation |
| --- | --- | --- | --- | --- |
| BMF2 | 3 | M | Diamond Blackfan Anemia | Heterozygous RPL11 |
| BMF7 | 3 | F | Diamond Blackfan Anemia | Heterozygous RPL35A |
| BMF25 | 3 | F | Fanconi Anemia | Biallelic FANCC |
| BMF35 | 4 | M | Fanconi Anemia | Biallelic BRCA2 |
| BMF9 | 8 | M | Fanconi Anemia | Biallelic FANCA |
| BMF23 | 14 | M | Shwachman-Diamond Syndrome | Compound Heterozygous SBDS |
| BMF41 | 4 | M | Shwachman-Diamond Syndrome | Homozygous SBDS |
| BMF45 | 8 | M | Shwachman-Diamond Syndrome | No identified mutation |
| BMF52 | 8 | M | Shwachman-Diamond Syndrome | No identified mutation |

To quantify the relationship between CDTA and age across the healthy lifespan, we plotted CDTA for all non-DC donors (**Figure 4C**). The relationship between CDTA and age across the full non-DC cohort (n=28, age range 4-75 years) follows a power law: CDTA = 9.33·(Age)^-0.7^ (r² = 0.59, p = 1.9×10^-6^, **Figure 4C**). This quantitative relationship, in which CDTA declines steeply in childhood and more gradually in adulthood, provides a reference curve against which individual CDTA values can be interpreted in future studies.

We next asked if this loss in CDTA in the DC cohort was biased towards either shorter or longer footprints. We determined the fraction of short footprints (populations I and II as defined in **Figure 2B**) and long footprints (population III as defined in **Figure 2C**) for Non-DC BMF and DC cohorts. We observed a significant decrease in short footprints and an increase in long footprints trending towards significance in the DC cohort (**Figure 4D**, **Supplementary Figure S3C**). This loss of shorter footprints suggests a loss of footprints consistent with TRF1/2 binding in DC. To ask if CDTA reflects shelterin binding, we performed CUT&RUN for TRF1 across seven PBMC samples for which we had matching CDTA measurements (DC, n=4; Pediatric healthy donor, n=3). We observe significant correlation between CDTA and fraction of telomeric fragments from TRF1 CUT&RUN (r^2^ = 0.66, p = 0.027, **Supplementary Figure S3D**). As a genomic control, we also were able to perform H2B CUT&RUN on six PBMC samples with matched CDTA measurements (DC, n=4, Pediatric healthy donor, n=2), and we did not observe any correlation between CDTA and fraction of telomeric fragments from H2B CUT&RUN (r^2^ = 0.062, p=0.634, **Supplementary Figure S3E**).

We and others have shown that cfDNA fragment length distributions can reveal levels of accessibility and transcription^12^. The enrichment of subnucleosomal footprints corresponding to nucleosome unwrapping correlates with genomic activity at gene promoters and enhancers. At a 1 kb window adjacent to telomeres and at control regions (large gene deserts with minimal chromatin features, see **Methods**), the fragment length distributions are dominated by the nucleosomal footprints ∼167 bp (**Figure 4E**). We do not observe the specific subnucleosome peaks characteristic of telomeric sequences in these regions, possibly because these sequences lack TRF1/2 binding. Notably, the frequency of subnucleosomal footprints is higher in the 1 kb subtelomeric window adjacent to telomeres compared to control regions (35-100 bp). Additionally, the dinucleosome peak is left-shifted at the subtelomeric window compared to control regions, reminiscent of increased transcriptional activity^38^. One possible reason for the shifts in cfDNA footprints within the 1 kb window adjacent to the telomere compared to control regions could be due to transcription that produces TERRA transcripts^48^. We asked if the subnucleosomal fragments are higher in the DC cohort due to dysfunctional telomeres. We calculated the log2 ratio of the frequency of footprints in 10 bp bins, comparing the 1 kb subtelomeric window to control regions for non-DC BMF and DC cohorts. We observed an increase in normalized frequency of fragments <100 bp (denoted by “**#**”, **Figure 4F**) and a decrease in frequency of fragments ∼330-400 bp in the DC cohort compared to non-DC BMF cohort (denoted by “**\***”, **Figure 4F**). Fragments <100 bp are consistent with footprints of unwrapped nucleosomes, whereas fragments 330-400 bp have footprints of dinucleosomes. We calculated subnucleosome enrichment (SE) using a formula that incorporates all fragment classes (see ***Methods*** for the formula). We normalized subtelomeric SE to that of control regions and compared the normalized SE of the DC cohort to the non-DC BMF and pediatric healthy cohort. All of the ten 1 kb windows (starting from the edge of the telomere towards the centromere) feature higher SE for the DC cohort compared to the non-DC BMF cohort and pediatric healthy cohort (**Figure 4G**). Subtelomeric regions are normally maintained in a silenced state that depends on intact telomere length and shelterin occupancy^49^. Our results suggests that the loss of telomeric footprints and the reduced telomere length correlate with changed chromatin landscape at the telomere, which is no longer conferring a silent heterochromatic state. This could result in increased accessibility and possibly transcription in the subtelomeric region as reflected by the increased cfDNA SE (**Figure 4G**). In summary, DC results in a decreased abundance of telomeric footprints in cfDNA, a decrease in short footprints at the telomeres suggestive of the loss of TRF1/2 binding, and a concomitant increase in cfDNA SE at subtelomeric regions suggestive of increased accessibility and/or transcription.

### Promoter subnucleosome enrichment (PSE) identifies DC-specific gene signatures from cfDNA

Given the strong cfDNA signature at telomeres and subtelomeres in patients with DC, we next investigated the effect of DC on cfDNA signatures in the rest of the genome. We calculated the promoter subnucleosome enrichment (PSE) for each gene and donor as a proxy for active transcription. We first identified the nucleosome position immediately downstream of the transcription start site (TSS) for each gene in each cohort (referred to as the “+1 nucleosome”). We then identified cfDNA fragments within 100 bp of the +1 nucleosome position of each gene to calculate the PSE for that gene for each donor. Thus, we obtained ∼16,000 PSE values across 31 donors. PSE exhibits a strong correlation with gene expression (**Supplementary Figure S4A)**. We performed principal component analysis (PCA) on the PSE matrix across donors and observed that the loadings of PC2 and PC3 separated the four cohorts (**Supplementary Figure S4B**). We observed a significant linear correlation between PC3 and chronological age (**Figure 5A**, r^2^ = 0.77, p = 9.5×10^-11^). PC2, on the other hand, displayed a significant linear correlation with CDTA (**Figure 5B**, R^2^ = 0.31, p = 0.001) and was significantly elevated in the PED cohort compared to DC and OLD cohorts (**Figure 5C**). Thus, PC2 and PC3 represent the scaling of genes that separate the contributions of chronological age and the telomere state to the observed promoter cfDNA signatures.

**Figure 5.**
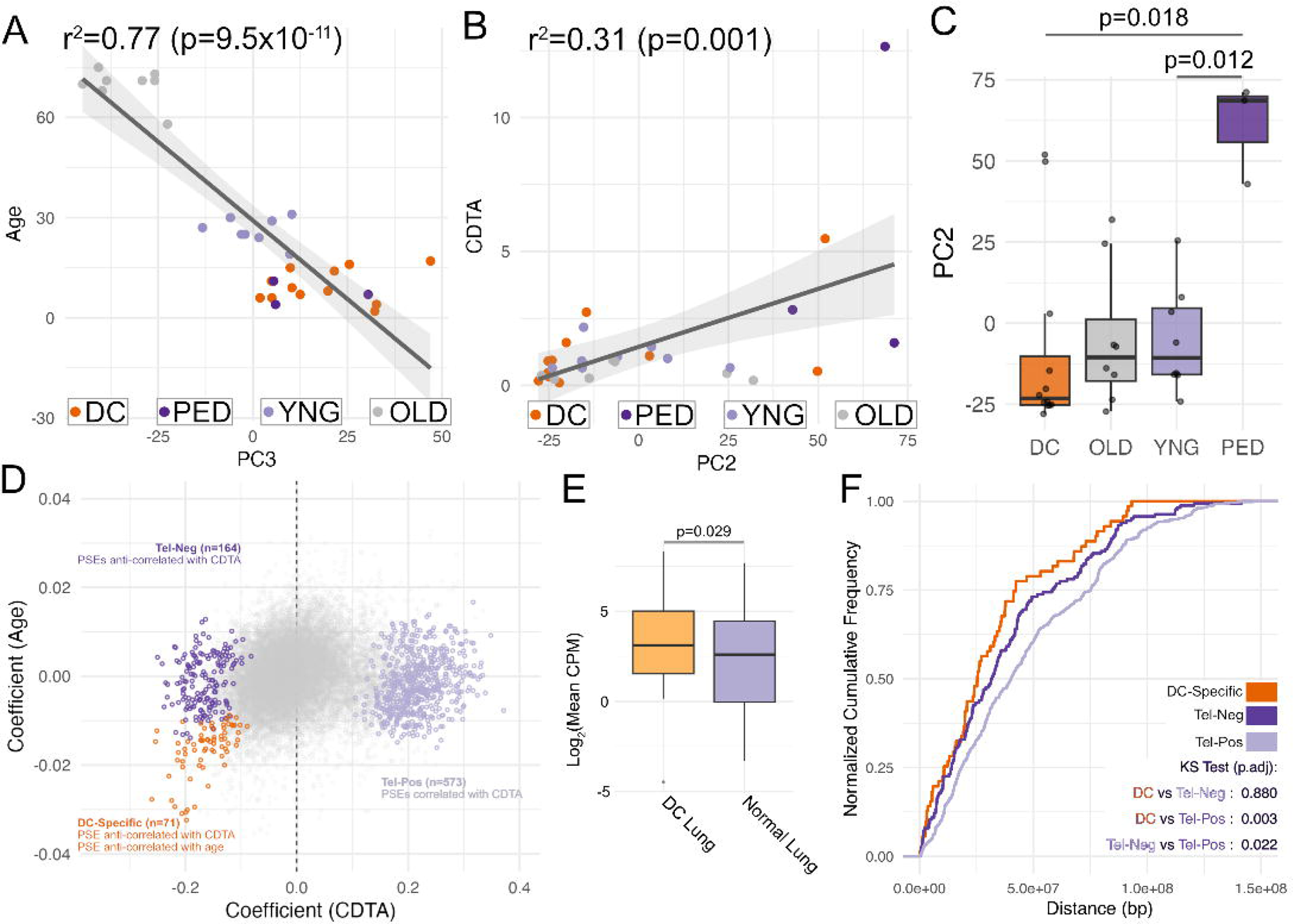
Promoter subnucleosome enrichment (PSE) identifies DC-specific gene signatures from cfDNA. **A**) Plot of Age vs. PC3 loadings, with line of best fit overlayed. The gray shaded region around the line of best fit represents the confidence interval (0.95) of the fit. Linear model: Age = a.PC3 + b; a=-0.94, 95% CI [-1.14, -0.75]; b=29.16, 95% CI [24.56,33.77]. P-value was calculated using F-test. F-statistic (1, 29) = 96.85. **B**) Plot of PC2 loadings vs. CDTA, with line of best fit overlayed. The gray shaded region around the line of best fit represents the confidence interval (0.95) of the fit. Linear model: CDTA = a.PC2 + b; a=0.04, 95% CI [0.02, 0.07]; b=1.44, 95% CI [0.71,2.16]. P-value was calculated using F-test. F-statistic (1, 29) = 13.14. **C**) PC2 loadings plotted for each cohort as points, overlayed by a boxplot. P-values were calculated using Wilcoxon Rank Sum test. **D**) Regression coefficients for CDTA (x-axis) and Age (y-axis) plotted for all genes for the linear model PSE = α.Age + β.CDTA. Genes with significant r^2^ and significant β are colored light purple (β>0, “Tel-Pos”), dark purple (β<0, “Tel-Neg”), and orange (β<0 and α>0, “DC-Specific”). **E**) Log2 of mean CPM of DC-specific genes for DC donors with fibrosis vs. healthy donors, from a lung RNA-seq dataset is shown as a boxplot. P-value was calculated using the Wilcoxon Rank Sum test with the alternative hypothesis that DC donors had higher expression compared to healthy donors. **F**) Cumulative distribution of distance of genes from telomere on the same chromosome arm for Tel-Pos, Tel-Neg, and DC-Specific group of genes defined in (**D**). P-values were calculated using Kolmogorov Smirnov test.

The PCA suggested that many genes’ PSE values could correlate with CDTA independent of age. To analyze this idea systematically, we fit a linear model for PSE against CDTA that regresses out age (PSE = α.Age + β.CDTA). Genes with r^2^ and β that are statistically significant would change significantly with changing CDTA, while controlling for age. We observed 164 genes with statistically significant negative β (“Tel-Neg”). These are genes whose PSE goes up as CDTA goes down. We also observed 573 genes with statistically significant positive β, genes whose expression positively correlates with CDTA (“Tel-Pos”). Finally, we observed 71 genes with statistically significant negative β and statistically significant positive α: genes whose PSE goes up with lower CDTA and younger age. These represent genes that have increased PSE specific to lower CDTA in the DC cohort which consists of donors 2-17 years old (**Figure 5D**).

We next asked whether the genes ranked as most dysregulated due to DC correlate with known phenotypes of DC. To obtain an independent measure of expression changes due to DC, we reanalyzed a published single-cell RNA-seq dataset of lung cells derived from healthy donors and those with lung fibrosis due to DC^50^. We calculated the mean counts per million (CPM) of the gene expression values across the DC and healthy cohorts. The genes identified to increase in PSE with decreasing CDTA and age, the “DC-specific” set feature significantly higher expression in the DC lung datasets compared to normal lung datasets (**Figure 5E**). Next, we asked if there is a specific positional bias in terms of distance to the nearest telomere for genes based on their change in cfDNA PSE relative to CDTA. When we plot the cumulative frequency of distance to telomere, DC-specific and Tel-Neg genes are significantly closer to the telomere compared to the Tel-Pos genes (**Figure 5F**). This positional bias in cfDNA PSE changes is consistent with derepression of normally silenced genes upon telomere shortening, as predicted by the phenomenon of Telomere Position Effect-Over Long Distances (TPE-OLD)^51^. In summary, we identified a promoter-specific cfDNA signature that changes with CDTA and age. The genes identified using this signature have correlated changes in expression in lung fibrosis and have a telomere distance-dependent pattern of change in PSE.

Finally, we asked if we could define a PSE signature for DC independent of CDTA and age. We compared PSE values across DC cohort (n=12, 2-17 years old) to the combined healthy cohort (n=19, 4-75 years old). We identified 183 genes with significantly higher PSE and 180 genes with significantly lower PSE in the DC cohort compared to the combined healthy cohort (**Supplementary Figure S4C**). For each donor, the average of PSE values for genes with Z-scores less than -2.5 were subtracted from the average of PSE values for genes with Z-scores greater than 2.5 to obtain a Δ score. The Δ score is higher for DC donors compared to healthy donors as expected (**Supplementary Figure S4D**). We then calculated Δ score for the three additional DC samples that were either collected at the same time or one year later, and also for the non-DC BMF samples. These datasets had not been used in the original Z-score calculations. These held out samples still showed clear separation between DC and Non-DC BMF samples, with DC samples having significantly higher scores (**Supplementary Figure S4E**). In summary, DC has a specific PSE signature that is distinct from healthy donors across age and donors with bone marrow failure due to non-DC diseases.

## DISCUSSION

The unique chromatin structure at the telomere, involving a combination of nucleosomes and shelterin complexes, is essential for chromosome end protection. Here, we show that human plasma cfDNA offers an unexpected means to obtain base-pair resolution footprints from the telomere, enabling us to propose models of telomeric chromatin structure that are consistent with observations from *in vitro* reconstitutions and cellular studies. We observe three predominant populations of telomeric footprints from short-read sequencing data, a subnucleosome species (population I, 49-66 bp), an unwrapped nucleosome species (population II, 115-132 bp), and nucleosomal species (>140 bp). Populations I and II arise from specific accessibility in the C-rich strand. The nucleosomal species could be resolved with long-read sequencing, showing a nucleosome repeat length of ∼145 bp.

Given the specific register of cleavage on the C-rich strand, it is tempting to speculate that population I arises due to shelterin footprints. Earlier studies have shown TRF1/2 binding up to 12 repeats *in vitro*, consistent with population I footprint sizes^52, 53^. Another possibility is that specific cleavages occur due to DNA accessibility resulting from TRF2 binding to columnar stacks of nucleosomes^54^. DNA sizes inferred from the mononucleosome-TRF1 cryo-EM structure match well with our short nucleosomal footprints (population II)^36^. Furthermore, if nucleosome wrapping is constrained based on TRF1 binding at the edge of the nucleosome, these nucleosomes are expected to have a specific positioning on the telomeric sequence due to TRF1’s binding register. As short nucleosomal footprints are included in the <139 bp footprints, they are enriched for the 5’ACCCTA and 5’CCCTAA registers as well, consistent with these footprints arising from unwrapped nucleosomes adjacent to a bound TRF1. Finally, the ∼145 bp nucleosome repeat length observed both on G- and C-rich strands is consistent with shorter nucleosome repeats seen in telomeric footprinting^39, 41^ and is consistent with repeat lengths observed with MNase footprinting of nucleosomes reconstituted on kilobases of telomeric repeat sequence^40^. Interestingly, the cryoEM structure from the same reconstitution showed a repeat length of 132 bp, with the α-N helix of H3 stacking against DNA that is part of the adjacent nucleosome. Our cfDNA footprints suggest a looser packing consistent with the reconstitutions in solution compared to the cryoEM structure in the same study^40^.

The two-category model of telomeric chromatin, one involving TRF1/2 binding adjacent to partially unwrapped nucleosomes, and the other involving compact nucleosome arrays with a ∼145 bp repeat length, is supported not only by our cfDNA footprint data but by convergent evidence from independent structural studies. The ∼123 bp short nucleosome footprint we observe in C-rich reads is quantitatively consistent with the ∼117 bp nucleosomal protection predicted from the cryo-EM structure of a TRF1-nucleosome complex, in which TRF1 binding at the nucleosome entry and exit sites unwraps approximately 1.5 turns of DNA on either side^36^. The ∼145 bp repeat length we observe for the phased nucleosome array is consistent with solution-state MNase footprinting of nucleosomes reconstituted on kilobases of telomeric repeat sequence, which revealed a repeat length distinct from, and longer than, the 132 bp repeat observed by cryo-EM of the same complex^40^. Additionally, in HeLa cells^41^ and rat liver nuclei^39^, MNase digestion followed by Southern blotting revealed shorter nucleosome repeat lengths at telomeres compared to the rest of the genome. Our cfDNA data thus bridges *in vitro* reconstitution and cellular studies: the footprint ensemble we observe in living humans is most parsimoniously explained by the same two structural states identified in these prior studies, suggesting that both states co-exist on telomeric chromatin *in vivo*. Direct visualization of these states in primary human cells by cryo-electron tomography or super-resolution imaging would be a powerful future direction to establish their relative proportions and any cell-type or disease-state dependence.

The abundance of the telomeric footprints decreases with age and due to DC. cfDNA represents DNA protected from nuclease cleavages, and CDTA may not represent telomere length, but instead the chromatin state of telomeres. The state of telomeres, rather than their length, is essential in inhibiting DNA damage repair and preventing senescence^42^. Thus, CDTA might be a complementary readout compared to telomere length, which is usually measured in lymphocytes. Because cfDNA is contributed by cellular turnover across multiple tissues, we speculate that, with altered physiology, CDTA would also capture telomeric footprints from non-hematopoietic sources. Future studies could identify non-hematopoietic contribution to CDTA, which might help uncover telomeric chromatin dynamics in diseased tissues. A direct correlation between individual CDTA values and matched telomere length measurements from the same donors would further validate CDTA as a telomere state readout. Such a correlation requires a cohort spanning a wide dynamic range of telomere lengths in the same individuals, a range that is not available in the current study, where Flow-FISH measurements were obtained only for DC donors, all of whom fall below the 10th percentile for telomere length by age. We note this as a limitation and a priority for future work, ideally using a cohort that includes healthy donors with a range of telomere lengths, DC donors with short and intermediate telomere lengths, and DC donors undergoing therapy.

We also observe significant enrichment for specific chromosome location for genes with changes in cfDNA PSE in DC patients. Subtelomeric regions feature higher normalized SE in DC patients compared to healthy donors, and genes whose promoter SE anticorrelate with CDTA are closer to the telomere compared to the genes whose promoter SE positively correlate with CDTA. Our results suggest long-range chromatin dysregulation emanating from the telomere due to DC. Telomere position effect was first demonstrated in yeast^55^ and subsequently in human cells^51, 56, 57^, where intact telomeres silence nearby genes over distances of up to 1 Mb. TPE-OLD extends this over longer distances via chromosome looping contacts between telomeres and interstitial telomeric sequences^51^. Our observation that genes with increased PSE in DC are preferentially closer to telomeres is consistent with de-repression of TPE-silenced genes upon telomere shortening: as telomeres erode in DC, the long-range silencing influence on proximal genes is relieved, resulting in their transcriptional activation. This interpretation is further supported by the fact that DC-specific genes show elevated expression in DC lung fibrosis RNA-seq data, consistent with active transcriptional upregulation rather than passive changes. Untangling which specific genes are de-repressed due to loss of telomere contacts versus other consequences of DC would require high-resolution chromatin contact maps comparing normal-length and shortened telomeres.

Regardless of the mechanism, the correlation between PC2 scaling and CDTA indicates that we have a specific cfDNA gene signature associated with DC. This observation suggests the possibility of identifying genes that are deregulated for each patient during each blood draw. DC is a heterogenous and heterochronous disease. The ability to detect organ-specific dysfunction in its early stages can lead to more effective treatment options for patients. The gene-based cfDNA analysis we have outlined has the potential to offer personalized physiological monitoring for DC patients, helping them navigate the disease more effectively.

## METHODS

### Human Sample Information

De-identified, healthy human plasma was purchased from Vitalant. De-identified samples for healthy pediatric donors, patients with Telomere Biology Disorder/Dyskeratosis Congenita, Fanconi anemia, Diamond-Blackfan anemia, and Shwachman-Diamond Syndrome were collected from consented patients at Children’s Hospital of Colorado with IRB approval (COMIRB 19-2913) (**Table 1**). Written informed consent was received prior to participation. Telomere length quantitated in unique cell populations identified through Flow FISH analysis (Repeat Diagnostics, North Vancouver, BC) and genetic mutation identification performed as part of standard of care diagnostic workup.

Peripheral blood plasma from samples were isolated from EDTA tubes by initial centrifugation at 1300g x 15 minutes (brake off, room temperature) followed by transfer of plasma to a second vial for centrifugation at 2200g x 5 minutes to separate the platelet-rich fraction from the plasma. Isolated plasma was then aliquoted and stored at -80C.

### UltraPrep cfDNA Isolation

cfDNA extraction was performed using the UltraPrep method ^58^. Briefly, human plasma (1 to 10 mL) was thawed from −80°C and spun at 21,000 rcf (relative centrifugal force), 4°C for 5 to 10 min to pellet any cell debris. The blood plasma supernatant was transferred to new tubes. 1 mL of blood plasma was transferred to a 5 mL conical tube, and 10 µL of proteinase K (20 mg/mL) was added to each donor sample. 650 µL of Digestion Buffer (5M guanidine thiocyanate (GITC), 25% Tween 20, 50 mM Tris pH 8.0, and 25 mM ethylenediaminetetraacetic acid (EDTA)) was added to the plasma and mixed well by pipetting. The samples were incubated at 56°C for 1 hour. 3.3 mL of Binding Buffer (3.5M GITC, 45% isopropanol, 2.5% Tween 20, 10 mM Tris pH 8.0, and 1 mM EDTA) was added to each sample. 40 µL of 400 nm SPHERO Silica Superparamagnetic beads (Spherotech) were added to each blood plasma sample and mixed well by vortexing. The bead-plasma slurry was incubated for 10 minutes at room temperature (RT). The tubes were placed on a magnetic rack, and when the supernatant was clear (∼ 5 minutes), it was discarded. The tubes were then removed from the magnetic rack, the beads were resuspended and washed with 500 µL of Wash Buffer 1 (3M GITC, 30% isopropanol, 5% Tween 20, 40 mM bis-tris pH 6.0, and 2 mM EDTA). The tubes were placed back on the magnetic rack, and once the slurry was clear, the supernatant was discarded. The beads were resuspended and washed with 500 µL of Wash Buffer 2 (50 mM Tris pH 8.0, 0.5 mM EDTA, and 80% ethanol). The tubes were placed back on the magnetic rack, and once the slurry was clear, the supernatant was discarded. The beads were resuspended with 100 µL of 100% EtOH. The new sample slurries were moved to 1.5 mL tubes. The tubes were then placed on a 1.5 mL magnetic rack, and when the slurry cleared, the supernatant was discarded. The tubes were spun down briefly and returned to the magnetic rack to remove the remaining EtOH. With the tube caps open, they were air dried at 37°C for 2 minutes in a bead bath. 30 µL of nuclease-free H2O was added to each sample and carefully mixed by pipetting. The tubes were incubated at RT for 5 minutes and returned to the magnetic rack. Once the supernatant was clear, the DNA was eluted into a new tube for each sample. Extracted DNA was quantified using the Invitrogen Qubit dsDNA HS Assay.

### Sequencing Libraries

cfDNA libraries were generated using the SRSLY PicoPlus Base Kit (cat*. CBS-K250B-96*). 1 to 10 ng of cfDNA per sample was combined with Nuclease-Free (NF) H2O and *SRSLY ss Enhancer* in a 0.5 mL Axygen PCR tube (*Fisher* Scientific cat. *14-222-290)* on ice. Following incubation on ice for 2 minutes, tubes were moved into a 0.5 mL PCR block for a heat shock at 98°C (lid 105°C) for 3 minutes to allow denaturation. The tubes were then incubated on ice for 2 minutes and the PCR block was set to 37°C (lid temp 45°C). A phosphorylation master mix was created by combining the SRSLY Master Mix, SRSLY Adapter A, and SRSLY Adapter B, all provided from the *SRSLY Base Kit*. To each sample, a phosphorylation master mix was added for a final volume of 50 μL. Once the tubes were mixed well via vortex, they were incubated for 1 hour in a PCR block at 37°C (lid temp 45°C). cfDNA was isolated after incubation using AMPure XP Beads (*Beckman Coulter* cat*. A63881*) at a ratio of 0.54x, mixed with 100% Isopropanol at a ratio of 0.24x, and NF-H2O. A 0.5 mL magnetic rack was used to capture and wash cfDNA with 80% EtOH. cfDNA-beads were suspended in 15 μL of NF-H2O, and cfDNA was eluted into new 0.5 mL Axygen PCR Tubes. Purified cfDNA libraries were amplified via PCR with a master SRSLY PCR Mix (provided in the base kit) and NF-H2O. Uniquely barcoded primers, P5 and P7, with a final concentration of 0.25 μM each, were added to each sample. Samples were initially incubated at 98°C (lid 105°C) for 3 minutes, followed by 10 cycles of amplification [98°C for 20 seconds, 65°C for 30 seconds, 72°C for 30 seconds], 72°C for 1 minute, then a final hold at 12°C. After PCR, cfDNA libraries were captured and washed with AMPure XP beads at a ratio of 0.95x. The final elution was in 20 μL 0.1X TE buffer and transferred to 1.5 mL Eppendorf LoBind tubes *(cat. 022431021*). Eluted libraries were quantified via Qubit (*dsDNA Quantitation, High Sensitivity* kit cat. *Q32854)* and quality checked using an Agilent Tapestation (*D1000* tape cat. *5067-5582*). Sequencing libraries were stored at -20°C until shipped for sequencing on the Novaseq. All libraries contained dual barcodes and were subjected to 150 bp paired-end sequencing.

### CUT&RUN

Frozen Peripheral Blood Mononuclear Cells (PBMCs) were quickly and completely thawed at 37°C. The required number of cells per sample (550,000 per reaction) were transferred to either a 1.5 mL tube or a 15 mL conical tube, then pelleted by centrifugation at 600 x g for 5 minutes at room temperature in a swinging-bucket centrifuge. The supernatant was carefully removed by pipetting and discarded. The cell pellet was then resuspended in 100 µL per reaction of cold NE Buffer (20 mM HEPES-KOH, 10 mM KCl, 0.1% Triton X-10, and 20% glyecerol) and incubated on ice for 10 minutes. 10 µL from each sample was removed to check nuclei integrity by microscopy. Samples were then centrifuged again at 600 x g for 5 minutes at room temperature in a swinging bucket centrifuge, the supernatant was discarded, and the cell pellet was resuspended in 100 µL per reaction of cold NE Buffer.

BioMag Plus Concanavalin A beads (ConA beads, Cat. 86057) at 11 µL per reaction were subject to two washes with Bead Activation Buffer (20 mM HEPES-KOH pH 7.9, 10 mM KCl, 1 mM CaCl2, and 1 mM MnCl2) at 100 µL per reaction. After the second wash, beads were resuspended in 11 µL per reaction of Bead Activation Buffer, homogenized by pipetting, and 10 µL of activated beads were added to one PCR tube per sample. Tubes were kept on ice until needed. Resuspended nuclei (in 100 µL volume) were added to each PCR tube containing 10 µL of activated beads, and the cell-bead slurry was homogenized well by pipetting and incubated for 10 minutes at room temperature.

During the bead-binding incubation, antibody master mixes are prepared for each target. For each antibody, the master mix was made by combining 50 µL of Antibody Buffer (20 mM HEPES-KOH pH 7.5, 150 mM NaCl, 0.5 mM Spermidine, Roche Complete Mini EDTA-free Protease inhibitor 1 tablet per 10 mL, 0.01%, Digitonin, and 2 mM EDTA) per sample with the appropriate volume of antibody. 1 µg of antibody was used for TRF1 (TRF1 antibody a gift from Dr. Joe Nassour) and 0.5 µg of antibody was used for H2B (Abcam Cat. ab1790). Tubes were placed on the magnetic rack until the slurry cleared, and the supernatant was removed and discarded. Tubes were removed from the rack, and the appropriate volume of antibody master mix was added to each sample. Samples were briefly vortexed to submerge the beads and homogenized by pipetting or flicking. Samples are then incubated overnight at 4°C on a nutator.

Following the overnight incubation, samples were briefly centrifuged, placed on the magnetic rack until clear, and the supernatant is removed and discarded. Keeping the tubes on the magnet, 200 µL of cold Digitonin Buffer (20 mM HEPES-KOH pH 7.5, 150 mM NaCl, 0.5 mM Spermidine, Roche Complete Mini EDTA-free Protease inhibitor 1 tablet per 10 mL, and 0.01% Digitonin) was added directly onto the beads, and the supernatant was immediately removed without disturbing beads. This wash was repeated once more for a total of two washes, with tubes kept on the magnet throughout.

Before removing the supernatant from the second Digitonin Buffer wash, the MNase master mix was prepared by combining 50 µL of Digitonin Buffer and 2.5 µL of pAG-MNase per sample (52.5 µL total per sample). After the second wash, supernatant was discarded, the tubes were removed from the magnet, and the 52.5 µL of MNase master mix was added to each reaction. Beads were resuspended by flicking the tubes and incubated for 10 minutes at room temperature on a nutator. During this incubation, the CaCl₂ master mix was prepared by combining 50 µL of Digitonin Buffer and 1 µL of 100 mM CaCl₂ per sample. Samples were placed back on the magnet, supernatant was removed, and two washes with 200 µL cold Digitonin Buffer were performed as described above. After the second wash, tubes were removed from the magnet, placed on ice, and 51 µL of the CaCl₂ master mix was added. Beads are resuspended by flicking, and samples were incubated for 2 hours at 4°C on a nutator.

To stop MNase digestion, 33 µL of stop buffer (340 mM NaCl, 20 mM EDTA, 4 mM EGTA, 50 µg/mL RNase A, and 50 µg/mL Glycogen, 0.5 ng of 0.5 ng/µL spike-in DNA per sample) was added to each sample, beads were resuspended by gentle vortex, and Proteinase K and SDS were added to each sample (8.4 µL of 10% SDS and 2.5 µL of 20 mg/mL Proteinase K per 84 µL supernatant volume). Samples were gently vortexed to mix and incubated for 10 minutes at 70°C in a thermocycler with the lid set to 75°C. Each sample was brought to 300 µL by adding 0.1X TE and 300 µL phenol-chloroform (1:1 ratio). Samples were stored at -20°C as a pause point until DNA purification was performed.

Samples were removed from -20°C and thawed at room temperature. Samples were mixed by inversion and centrifuged at 20,000xg for 5 minutes at room temperature. The upper aqueous phase was carefully transferred to a new 1.5 mL tube. A second extraction was performed by adding phenol-chloroform in a 1:1 ratio to the collected aqueous phase, mixing by inversion, and centrifuging again. The upper phase was again carefully transferred to a new tube. A third and final extraction was performed by adding chloroform in a 1:1 ratio to the collected aqueous phase, mixing by inversion, and centrifuging as above. The upper aqueous phase was transferred to a new tube. The final aqueous phase was subjected to a 0.5X AMPure bead cleanup. Beads and sample were combined at a 0.5X bead-to-sample ratio, mixed, and incubated for 10 minutes at room temperature. Samples were placed on the magnetic rack to separate, and the supernatant (unbound fraction, which contains shorter fragments) was transferred to new tubes for a subsequent 2X AMPure bead cleanup. The bound fraction was retained separately. For the 2X AMPure cleanup, beads and sample were combined at a 2X bead-to-sample ratio, mixed, and incubated for 10 minutes at room temperature. Samples were placed on the magnetic rack, and the supernatant was discarded. Beads were washed twice with 500 µL of freshly prepared 80% ethanol, returning the tubes to the magnet and discarding the supernatant between each wash. Beads were then air-dried outside of the magnetic rack for 3-5 minutes and resuspended in 30 µL of nuclease-free water to elute. After a 15-minute incubation at room temperature, tubes were placed on the magnetic rack, and the eluate is transferred to new 0.5 mL low-bind tubes. DNA concentration was quantified using the Qubit fluorometer and sequencing libraries were generated and paired-end sequencing performed in the same manner as for cfDNA samples.

### PCR enrichment of telomere sequences from SRSLY cfDNA libraries

cfDNA libraries until the PCR amplification step were generated as described above. To enrich for telomeric sequences for long-read sequencing, the libraries were amplified using CloneAmp HiFi PCR premix, which accurately amplifies long amplicons. Either standard P5/P7 primers or primers that recognize the junction between Illumina adapters and telomeric repeats were used for amplification (primer sequences below). We used three primer sets:

1. P5 standard primer sequence and P7 standard primer sequence
2. P5 standard primer sequence and P7 + telomere repeat sequence
3. P7 standard primer sequence and P5 + telomere repeat sequence

The PCR conditions were (lid temp 105°C), 98°C 10 seconds, 52°C 15 seconds, 72°C 5 seconds, repeated for 12 total cycles, and a 10°C holding temperature. The cfDNA PCR product was purified using a 2X AMPure bead clean. Eluted libraries were quantified using Qubit (dsDNA Quantitation, High Sensitivity kit cat. Q32854) and quality-checked using an Agilent Tapestation (D1000 tape cat. 5067-5582). Sequencing libraries were stored at -20°C until utilized for Nanopore sequencing.

P5 standard primer sequence: 5’ TTTCCCTACACGACGCTCTT 3’

P7 standard primer sequence: 5’ GTTCAGACGTGTGCTCTTCC 3’

P5 + telomere repeat sequence: 5’ TCTTCCGATCTCCCTAACCCTA 3’

P7 + telomere repeat sequence: 5’ CTCTTCCGATCTCCCTAACCC 3’

### Oxford Nanopore Long Read Sequencing

The Nanopore Ligation Sequencing DNA SQK-LSK 110 and SQK-LSK 114 kits were used to generate cfDNA Nanopore sequencing libraries, which were sequenced on R9.4.1 MinION and R10.4.1 PromethION flow cells, respectively. The basecalling models used were dna_r9.4.1_e8_sup@v3.3 and dna_r10.4.1_e8.2_400bps_sup@v4.2.0.

### Single-end analysis

We first performed analysis using only the R1 read, due to the ambiguities involved in aligning paired-end reads to repetitive DNA. Adapter sequences were trimmed, and reads at least 18 bp long were selected using Cutadapt (version 5.2)^59^. The number of reads with GC content between 48 and 52% was calculated. For footprint analysis, reads were further trimmed to 140 bp. This ensured that we could confidently assign footprint lengths of 18 to 139 bp. The actual lengths of 140 bp reads are unknown by Illumina sequencing. Footprints close to nucleosomal sizes (∼147 bp) would have adapter sequences too short to be identified; therefore, to achieve the highest confidence in footprint lengths, we trimmed reads with lengths greater than 140 bp down to 140 bp. For the analysis shown in Figure 1, we then used a custom Python script (input_6mer.filter.py) to identify reads composed of 6-mer repeats. For example, for CCCCCA, we scan each read in 1-bp steps for all circularly permuted versions of CCCCCA. The read is assigned to CCCCCA if all 6-mers generated from the read have edit distances equal to 0.

For determining coverage of 6-mers in the reference genome, the RepeatMasker ^60^ output for the hs1 v2 genome was mined for every 6-mer entry. Then the bases covered by each entry were added, combining the base counts for circularly permuted versions of the same 6-mers, similar to the analysis performed for cfDNA.

For analysis of reads with telomeric repeats (Figures 2 and 4), we used a custom Python script (filter_kmer_v2.py) to filter for reads with telomeric repeats. 6-mers were generated by scanning reads in one bp steps, and the lowest edit distance between the generated 6-mer and all rotational permutations of the G-rich and C-rich telomeric repeat was determined. If 100% of the 6-mers generated from the read had edit distance less than or equal to 1, that read was selected as made of telomeric repeat sequence. The number of reads made of the telomeric repeat sequence was used for the calculation of cell-free DNA telomeric abundance (CDTA):

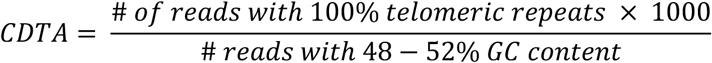

### Subtelomeric and Promoter Subnucleosome Enrichment

Illumina cfDNA datasets were processed with Cutadapt (version 5.2)^59^ in paired-end mode to remove Illumina adapters. Only reads at least 35 bp after removal of adapters were retained. Reads were trimmed down to a maximum of 130 bp using Cutadapt. Trimmed reads were aligned to the human reference genome (T2T CHM13v2.0/hs1) using bowtie2 (version 2.5.4, with parameters “--local --very-sensitive-local --no-unal --no-mixed --no-discordant -I 10 -X 500”). Samtools^61^ and bedtools^62^ were used for processing aligned reads from SAM to BED files. Duplicate reads were discarded for further analysis if the reads had the same start and end coordinates.

Based on the reference genome telomere coordinates, subtelomeric windows of 1 kb length were defined starting from the edge of the telomeres. Coordinates for the gene deserts, which are expected to have minimal chromatin dynamics are as follows: chr4:114003078-115431124, chr5:76212901-76548648, chr2:147414195-148264716, chr2:155320669-155917930, chr2:154926419-155132433, and chr13:82000000-85000000. The unique cfDNA reads mapping to subtelomeric windows and α-satellite regions were identified using bedtools intersect. These fragments were used to generate fragment length distributions in 10 bp bins shown in Figure 4E and F. Subnucleosome enrichment was calculated using the following formula:

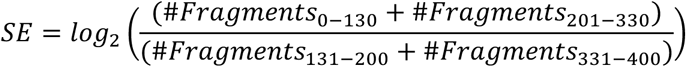

The subtelomeric SE was normalized for each sample by subtracting the average SE value across the gene deserts.

To calculate SE values for genes (promoter subnucleosome enrichment, PSE), first the +1 nucleosome for each gene was defined individually for DC, YNG, and OLD cohorts. Nucleosome positions were called using a custom Perl script (https://github.com/srinivasramachandran/CallNucleosomes) and the +1 nucleosome position was defined as the first nucleosome position downstream of the TSS that is between the TSS and the TSS + 300 bp. For each donor, for each gene, reads mapping within 100 bp of the +1 position of that gene were used to calculate SE for that gene.

### Nanopore Long-Read Datasets

Base-calling was performed with Dorado. Since P5/P7 adapters were ligated before ONT library preparation, we could use them as anchors to precisely define read lengths. Hence, we wrote custom python scripts (v3.adapter_trim_write_NoTrim.py, v2.adapter_trim_P7TelP5Pln.py, v2.adapter_trim_P5TelV2P7Pln.py) to trim P5 and P7 adapters from ONT reads. After trimming, filter_kmer_v2.py was used to filter for reads with telomeric repeats. Reads were also aligned to the CHM13 T2T hs1 v2 human reference genome using minimap2.

### Other Analyses

The fibrosis RNA-seq dataset was obtained from GEO (Accession: GSE83501, file: GSE83501_human_FPKM_matrix.txt.gz). The matrix from the GEO accession contains expression values in Fragments Per Kilobase of transcript Per Million mapped reads (FPKM) for single cells sequenced from the lungs of different patient cohorts. We selected the healthy and DC cohorts for our analysis. We first converted FPKM of each gene to transcripts per million (TPM) using the following formula:

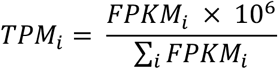

where TPM_i_ denotes TPM of gene i, and FPKM_i_ denotes FPKM of gene i. Then, the mean and SEM of TPM values for each gene across each cohort were calculated. Then, a Z-score was calculated for each gene as follows:

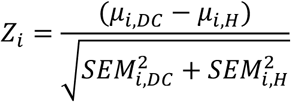

Where Z_i_ is the Z-score for gene *i,* μ_i,DC_ is the mean expression of gene *i* across the DC cohort, μ_i,H_ is the mean expression of gene *i* across the healthy cohort, S.E.M_i,DC_ is the standard error of mean of gene *i* across the DC cohort, and S.E.M_i,H_ is the standard error of mean of gene *i* across the healthy cohort. The Z-score captures changes in gene expression in fibrotic lung cells from DC donors compared to non-fibrotic lung cells from healthy donors.

The “bedtools closest” command was used to calculate the distance of genes from the telomere. A BED file containing transcription start sites for each gene and a BED file with telomere coordinates were used as inputs. This distance was used to bin genes.

To bin genes based on change in their PSE in the DC cohort compared to the healthy young cohort, we calculated the Z-score of PSE of each gene, similar to the approach used for RNA TPM values above.

## DATA AVAILABILITY

Sequence data that support the findings of this study have been deposited in in dbGaP with the accession code phs004409.v1.p1 (https://dbgap.ncbi.nlm.nih.gov/). Source Data are provided with this paper.

## CODE AVAILABILITY

All custom code generated for analysis presented in this study have been deposited in GitHub, in the repository: https://github.com/srinivasramachandran/6merAndTelomere

## ACKNOWLEDGEMENTS

We would like to thank Dr. Sujatha Jagannathan and Dr. Joe Nassour for critical reading of this manuscript. We would like to acknowledge support from NIH grant R35GM156411 (S. R.), NSF Award #2330283 (L. K. W.), American Cancer Society RSG-22-026-01 (S.R.), the Pew Charitable Trusts and the Alexander and Margaret Stewart Trust (S.R.), the Boettcher Foundation’s Webb-Waring Biomedical Research Program (C.F.) and Garborini Memorial Fund (C.F.).

## AUTHOR CONTRIBUTIONS

Brandon Buck: Conceptualization, Methodology, Investigation, Data Curation, Writing (Original Draft). Abigail Weirich: Methodology, Validation, Investigation, Data Curation. Stephanie Eramo: Methodology, Validation, Investigation. Rachel Lopez: Investigation. Laura K. White: Methodology, Writing (Review & Editing). Peter Kabos: Methodology, Writing (Review & Editing). Craig Forester: Conceptualization, Methodology, Validation, Investigation, Resources, Data Curation, Writing (Review & Editing), Supervision, Project Administration, and Funding Acquisition. Srinivas Ramachandran: Conceptualization, Methodology, Software, Validation, Formal Analysis, Resources, Data Curation, Writing (Original Draft), Visualization, Supervision, Project Administration, and Funding Acquisition.

