## Supplementary Figures for "Fragmentation Patterns of Human Telomeric Chromatin in Plasma cfDNA"

### Supplementary Figure 1

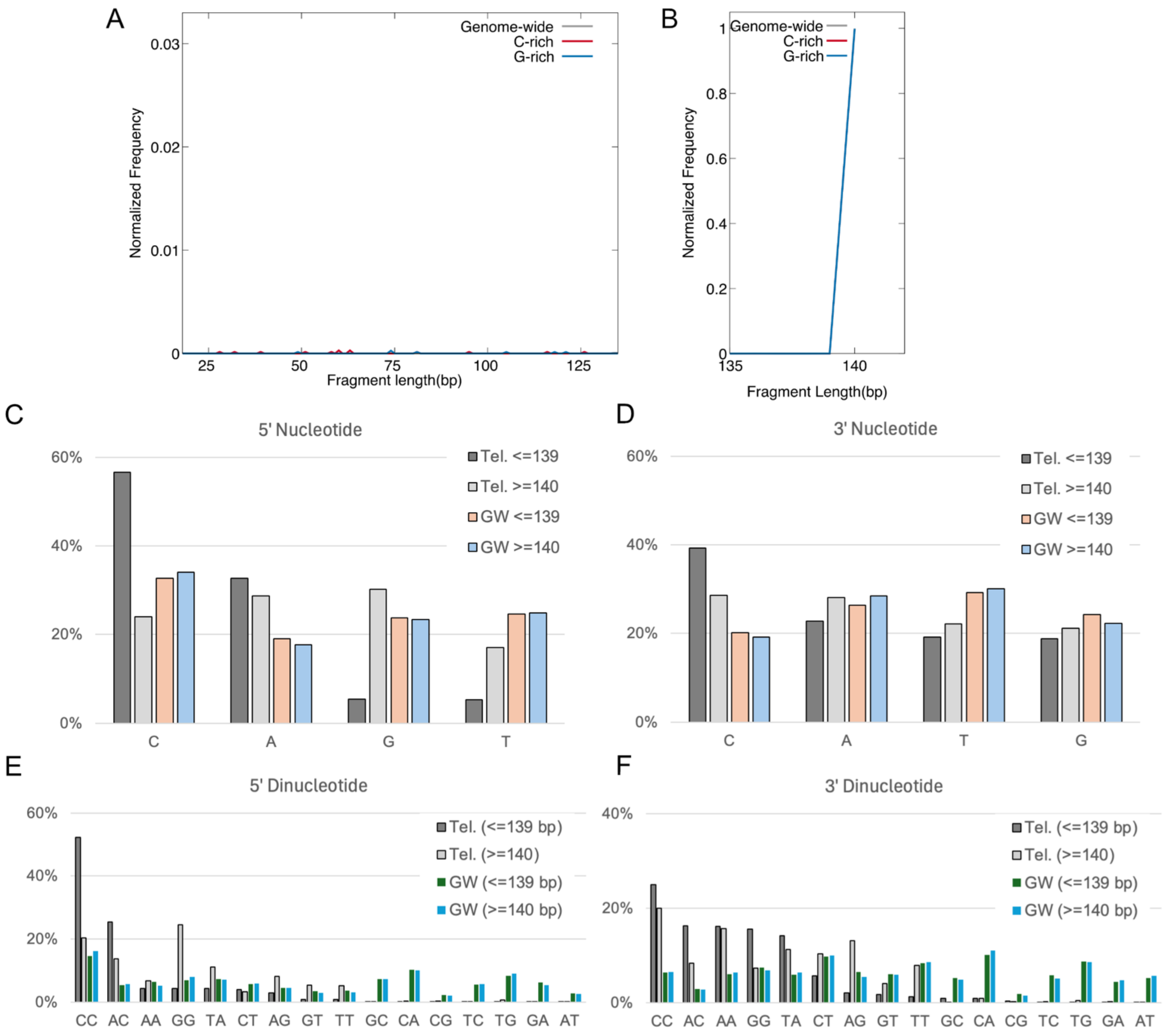


**Supplementary Figure S1. Double strand library and nucleotide preferences. A**) Fragment length distribution of the telomeric cfDNA in the range 18-135 bp for published datasets generated with double-strand library protocol. **B**) Same as (**A**) for fragments in the range 135-140 bp. **C**) 5’ nucleotide frequency for fragments defined as telomeric (“Tel.”) and genome-wide (“GW”) for fragments <=139 bp and >=140 bp in length. **D**) Same as (**C**) for the 3’ nucleotide. **E**) Same as (**C**) for 5’ dinucleotide. **F**) Same as (**C**) for 3’ dinucleotide.

**Supplementary Figure 2**

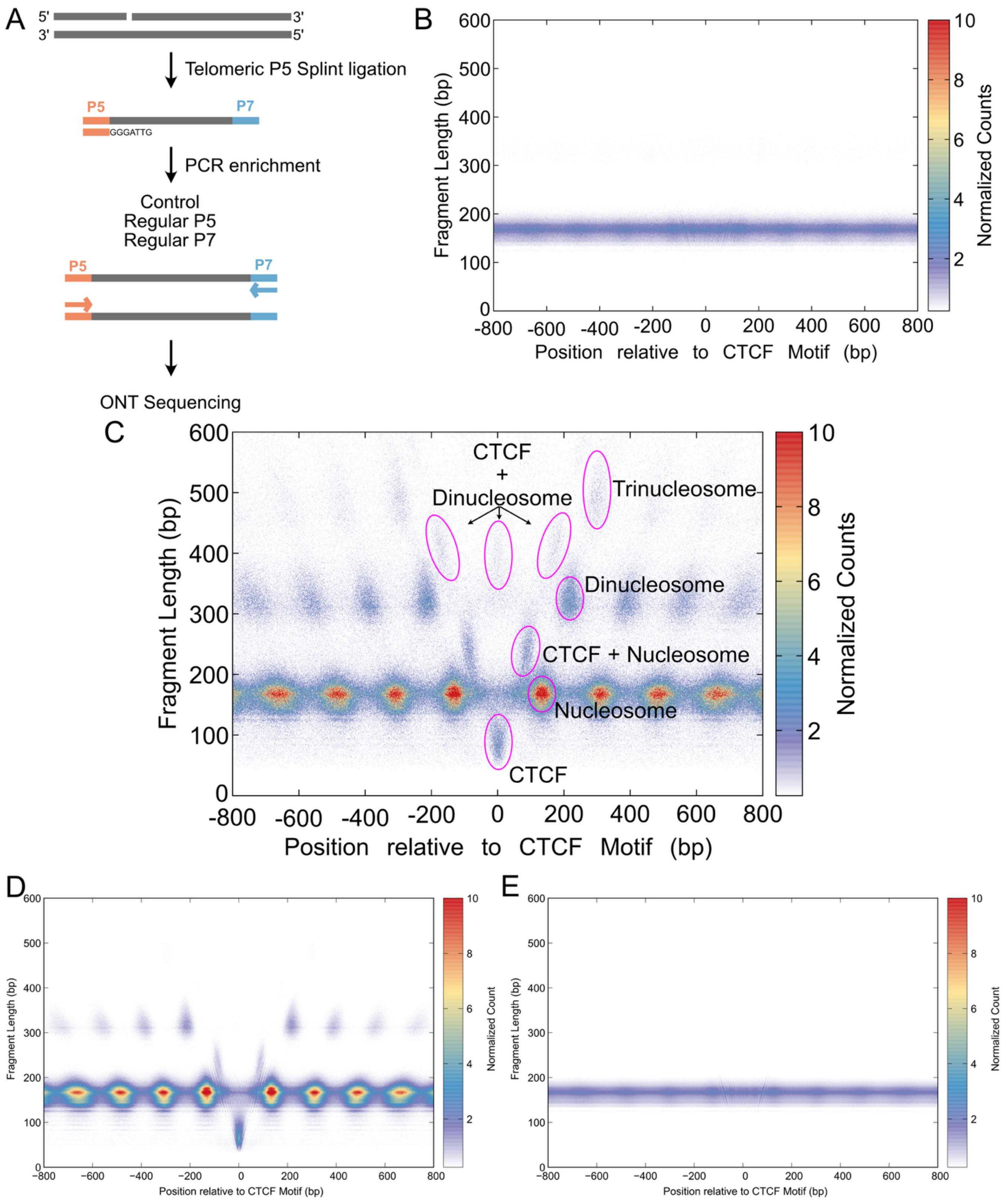


**Supplementary Figure S2. Long-read sequencing of cfDNA.** A) Schematic of library strategy used to enrich for telomeric sequences from cfDNA while preserving fragment ends, by using a specific splint sequence that anneals to telomeric repats. **B**) Fragment midpoint vs. length plot ("V plot") at CTCF motifs that are depleted for short footprints in cfDNA generated using nanopore sequencing data. **C**) Annotated fragment midpoint vs. length plot ("V plot") at CTCF motifs that are enriched for short footprints in cfDNA, denoting the inferred species for each population of footprints. **D**) V plot at CTCF motifs that are enriched for short footprints in cfDNA, generated from short-read sequencing data. **E**) Same as (**D**) at CTCF motifs that are depleted for short footprints in cfDNA.

**Supplementary Figure 3**

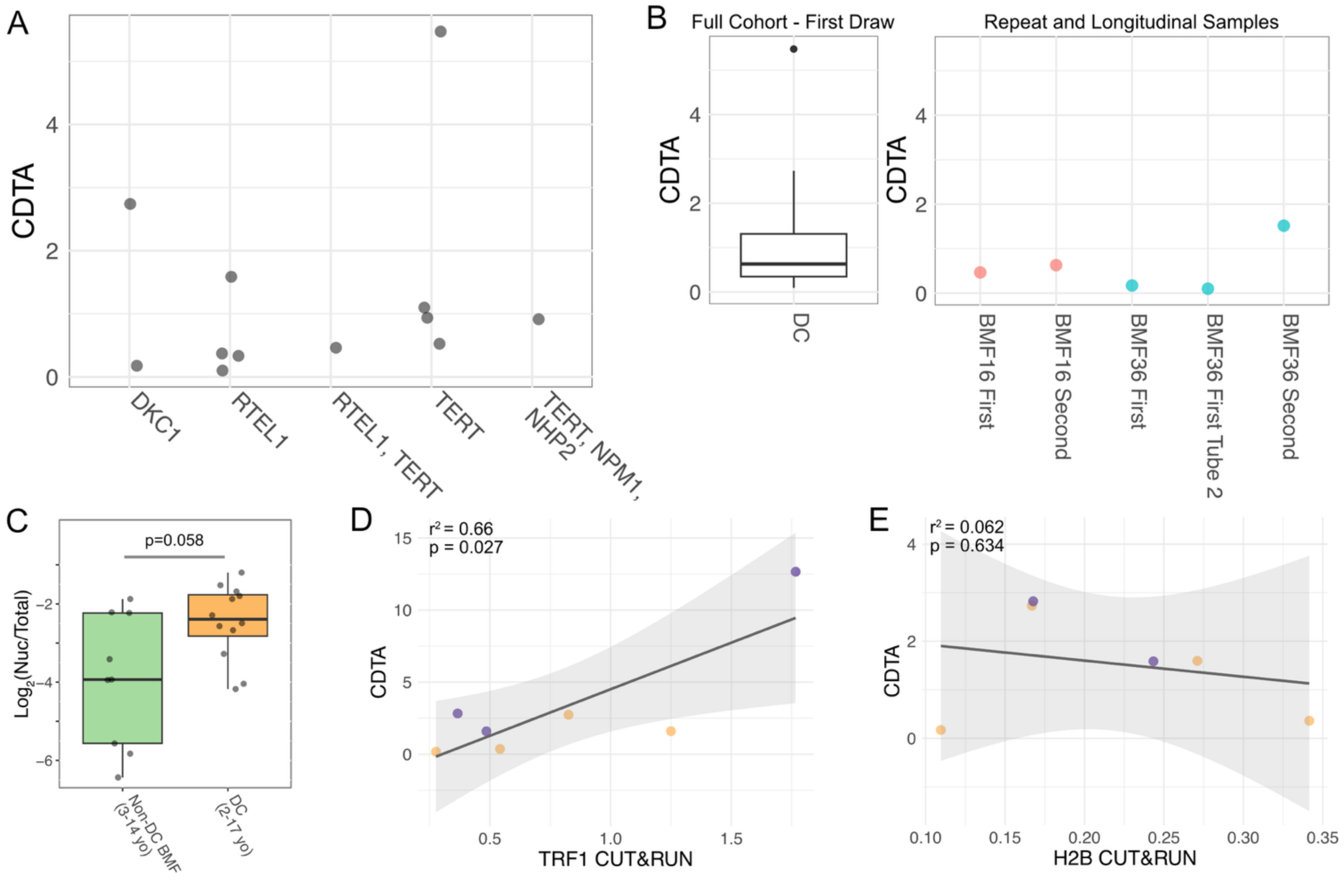


**Supplementary Figure S3. Characterization of telomeric cfDNA of DC cohort. A**) Cell-free DNA telomeric abundance (CDTA) for Dyskeratosis Congenita (DC) donors stratified by mutation status. No mutation-specific trend was observed. **B**) Comparison of CDTA for DC donors determined from first blood draw (n=12, left) and for those donors with multiple blood draws (right). BMF16 donor had two blood draws one year apart. BMF36 donor had two tubes of plasma from first draw which were processed on independent days but yielded highly similar CDTA values, and a blood draw one year later, which also yielded CDTA value within range of the first draw of the whole DC cohort. **C**) The 140 bp nucleosomal footprints were quantified for Non-DC BMF and DC donors, and the distribution across the cohorts is shown as a boxplot. P-values was calculated using the Wilcoxon Rank Sum test. **D**) Comparison of CDTA and abundance of telomeric fragments from TRF1 CUT&RUN performed on matched PBMC samples. Orange denotes DC samples and purple pediatric healthy samples. **E**) Same as (**D**) for H2B. P-values were calculated using the F-test.

**Supplementary Figure 4**

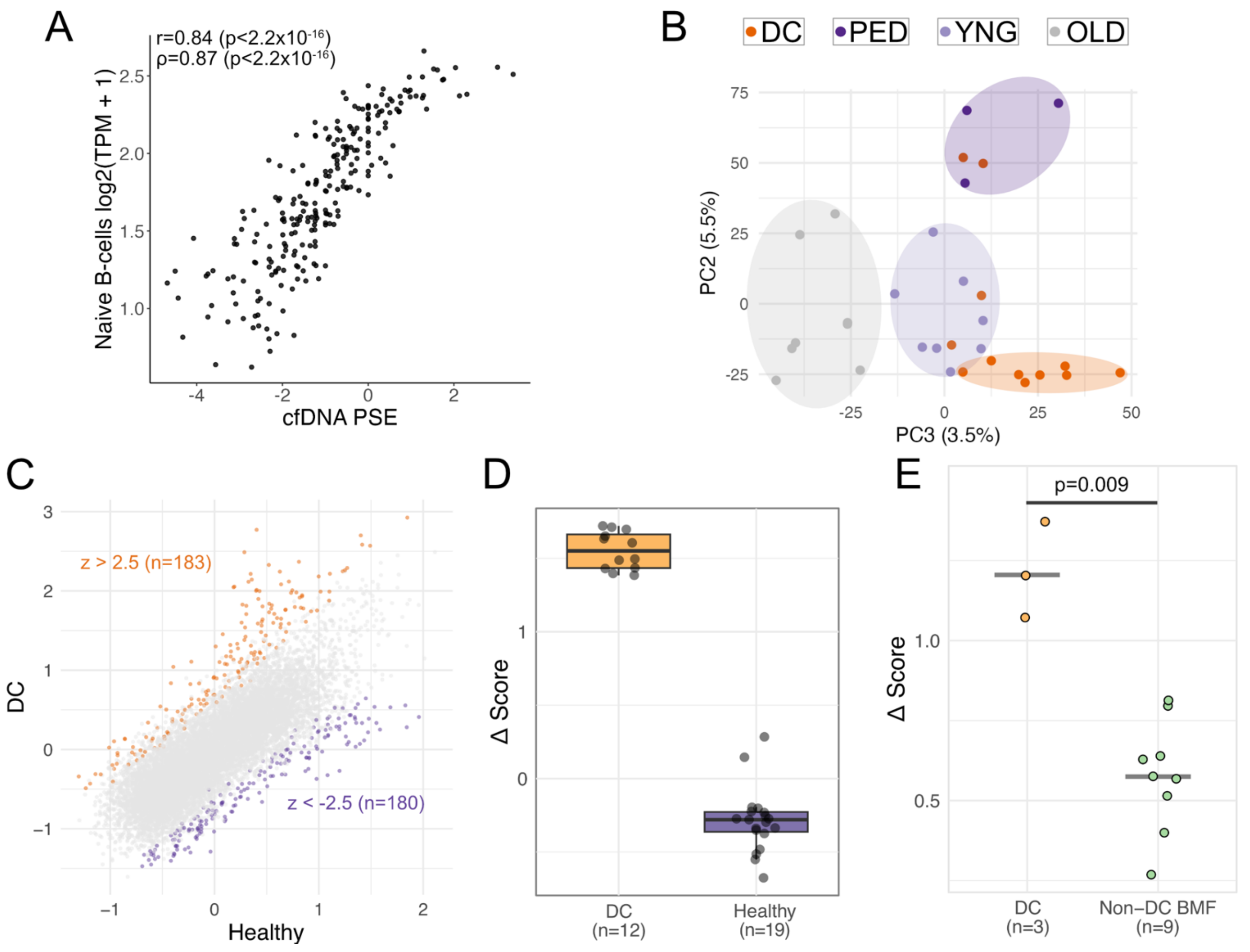


**Supplementary Figure S4. Promoter subnucleosome enrichment analysis. A**) Genes were sorted by their cfDNA PSE for a representative healthy donor, and the reads were combined for 50 genes in each bin to calculate the cfDNA PSE. The cfDNA PSE is plotted against a published RNA-seq dataset [82] from naïve B-cells. RNA-seq was also averaged within the same bins as cfDNA. Pearson r and Spearman ρ values for the correlation between cfDNA and RNA-seq are shown. **B**) Principal Component Analysis was performed on the cfDNA promoter subnucleosomal enrichment score matrix across healthy pediatric (n=3), healthy young (n=8), healthy old (n=8), and Dyskeratosis Congenita (DC) donors (n=12). PC2 loadings vs. PC3 loadings are plotted with shaded ovals delineating the cohorts. **C**) Average PSE across DC donors (n=12) versus all healthy donors combined (n=19) is plotted. The genes with Z-scores greater than 2.5 are highlighted in orange, whereas genes with Z-scores less than -2.5 are highlighted in purple. **D**) For each donor, the average of PSE values for genes with Z-scores less than -2.5 (identified in (**C**)) were subtracted from the average of PSE values for genes with Z-scores greater than 2.5 (identified in (**C**)) to obtain a Δ score. The Δ score is higher for DC donors compared to healthy donors as expected.

**E**) The Δ score was calculated for the repeat samples of DC (n=3) and Non-DC BMF samples (n=9), which were not used in the original Z-score calculations. These held-out samples still show a significant separation between DC and Non-DC BMF samples, with the DC samples having a higher Δ score. P-values was calculated using the Wilcoxon Rank Sum test.
